# Zero-shot automated insulin delivery for type 1 diabetes via dynamic physiology-aware reinforcement learning

**DOI:** 10.64898/2026.05.25.727637

**Authors:** Junyoung Yoo, Vega Pradana Rachim, Yein Lee, Jaeyeon Lee, Sung-Min Park

## Abstract

Insulin therapy in type 1 diabetes requires constant dose adjustment based on blood glucose, meals, physiological states, and physical activity. This demanding self-management imposes a substantial burden and increases dosing-error risk, underscoring the need for automated insulin delivery (AID) systems that reduce user intervention. However, many current systems depend on fixed, individualized parameters and may not fully adapt to rapid or unobserved physiological changes. We developed the Dynamic Physiology-Aware Reinforcement learning Controller (DPARC), a zero-shot insulin optimizer that infers latent physiological dynamics from recent continuous glucose monitoring (CGM) and insulin-delivery history without prior personalization, carbohydrate announcements, or preset subject-specific parameters. DPARC uses a rolling 24-hour CGM and insulin-history window, but closed-loop operation can begin after 1 hour of observed data by initializing unobserved history with neutral normalized padding and progressively replacing it with observations. In silico, a single frozen DPARC policy adapted within 1 hour, improved time in range compared with a total daily insulin-conditioned reinforcement learning baseline, and approached the upper-bound performance of a fully personalized model under stochastic unannounced meals with randomized timing, carbohydrate amounts, absorption variability, and meal skipping. In supervised porcine studies under unannounced meals, DPARC maintained high time in range without manual configuration, supporting large-animal feasibility while prospective human evaluation is needed before clinical efficacy can be established. Learned latent representations correlated with physiological markers including insulin sensitivity and plasma insulin concentration, supporting physiological alignment and explanatory anchors. Collectively, these findings support DPARC as a preclinical proof-of-concept zero-shot AID framework for future supervised human evaluation.

## INTRODUCTION

Type 1 diabetes (T1D) is a chronic metabolic disorder caused by autoimmune destruction of pancreatic β-cells, resulting in absolute insulin deficiency that necessitates lifelong external insulin administration (*1*). Despite advances in continuous glucose monitoring (CGM), insulin pumps, and ultra-rapid-acting insulin (*2*), a substantial proportion of individuals with diabetes continue to experience recurrent episodes of hypoglycemia and hyperglycemia due to significant glucose variability (*3*). This variability arises from the interplay of exogenous factors such as meals and exercise, which directly alter glucose levels, and endogenous factors such as circadian rhythms and stress, which modulate insulin resistance (*4*). Consequently, people with diabetes face glycemic extremes, where persistent hyperglycemia increases the risk of long-term complications including retinopathy, nephropathy, and cardiovascular disease (*5*), while acute hypoglycemia poses immediate life-threatening risks (*6*). Given the chronic nature of diabetes and the need for rapid, continuous responses to fluctuating glucose levels, there is a pressing demand for fully automated insulin delivery systems that minimize the burden and potential errors associated with manual input (*7*).

To address these clinical needs, automated insulin delivery (AID) systems integrating CGM and insulin pumps in closed-loop configurations have been developed. These systems have progressed from early low glucose suspend (LGS) designs to hybrid closed-loop (HCL) and fully closed-loop (FCL) systems, with each generation reducing user intervention (*8*). Initial LGS systems suspended insulin only during hypoglycemic episodes, whereas HCL systems enabled automated basal insulin adjustments (*9, 10*). More recently, advanced hybrid closed-loop (AHCL) systems (*11*) have simplified meal announcements (e.g., low/medium/large) and further enhanced usability, moving closer to the vision of FCL systems (*12*) capable of achieving fully automated glucose regulation without any manual input.

However, current AHCL systems and FCL prototypes remain constrained by their reliance on predefined individual-specific parameters. These systems require manual configuration of personalization parameters, including total daily insulin (TDI), carbohydrate ratio (CR), and insulin sensitivity factor (ISF) for optimal insulin dosing (*13–15*). The accuracy of these parameters is critical to system performance but difficult to establish precisely. Studies have demonstrated that AHCL and FCL systems based on fixed parameters are vulnerable to estimation errors, which can exacerbate glucose variability and compromise glycemic control (*14, 15*). More importantly, fixed-parameter approaches cannot account for physiological fluctuations occurring on hourly, daily, or weekly timescales. Insulin sensitivity fluctuates significantly due to infection, stress, hormonal shifts, and physical activity, yet current systems cannot detect or adapt to these changes in real time (*16*). These limitations underscore the need for adaptive control strategies (*17*), as optimizing insulin dosing requires dynamic, real-time assessment of changing physiological states rather than static feedback control. Traditional model predictive control and proportional-integral-derivative controllers rely on predefined mathematical models, restricting their ability to address individual variability that is inherently difficult to capture (*18*).

Reinforcement learning (RL) offers a promising solution for adaptive insulin control because it learns directly from interaction and adapts to nonstationary conditions without requiring fixed personalization parameters or mechanistic models (*19, 20*). In this framework, in vivo glucose–insulin dynamics constitute the environment, CGM-derived histories define the state space, and insulin delivery actions form the control space. The reward function prioritizes time in range (TIR) while penalizing hypoglycemia, hyperglycemia, and abrupt excursions in different weights. Recent studies have shown that RL-based controllers can outperform conventional algorithms across diverse virtual cohorts and enhance long-horizon stability (*18, 21–24*), reflecting the benefit of optimizing long-term rewards that balance immediate correction with sustained glycemic stability (*24*). However, the trial-and-error nature of RL introduces safety risks (*25*). Unstable policies may emerge during early training, potentially leading to dangerous glycemic events, making direct online learning in people with diabetes unsafe. Therefore, RL-based controllers require extensive training in simulated environments before real-world deployment to ensure safety (*26, 27*). While this simulation-based training approach addresses safety concerns, it introduces a major limitation: the simulation-to-reality (sim-to-real) gap. This gap arises from distributional mismatches between simulator training domains and real deployment conditions, as simulators capture only limited historical cohorts. Consequently, policies trained in silico must extrapolate to new users and evolving physiological states, lifestyles, disease trajectories, and drug responses, creating a temporal mismatch between training and deployment without guaranteed real-world performance (*28*).

To address the aforementioned limitations, we propose a context-free adaptive RL approach that eliminates explicit reliance on personalization parameters and directly addresses temporal mismatch between pre-trained simulators and dynamic real-world physiology. The key idea is to pre-train the policy using domain randomization (*29*), exposing it to a wide range of physiological variability observed in diabetes. During deployment, the controller performs continual adaptation by inferring the user’s current physiological state from real-time CGM data, thereby aligning policy behavior with evolving physiology without requiring any additional training. This strategy is inspired by history-conditioned policies in robotics, where unobservable conditions such as terrain or friction are inferred from sensor histories (*30, 31*). Building on this concept, we developed the Dynamic Physiology-Aware RL Controller (DPARC), trained entirely in simulation to autonomously infer latent physiological features in real time. DPARC employs a temporal encoder that operates on a rolling 24-h CGM and insulin-history window, producing latent representations that reflect dynamic physiological factors such as insulin sensitivity and plasma insulin concentration. These features enable personalized insulin dosing without fixed parameterization (*32, 33*). Notably, although the encoder architecture uses a 24-h rolling window, DPARC can start closed-loop operation after approximately 1 h of observed CGM and insulin data by initializing the unobserved history window and progressively replacing it with observed data, without manual personalization or subject-specific parameter tuning.

To demonstrate DPARC’s practicality, we assessed performance in both in silico and multi-species preclinical models (Table 1). A single policy trained in a UVA/Padova T1DMS-based simulation environment demonstrated zero-shot generalization to independent virtual cohorts, and the same frozen policy was transferred to rodent and swine models of T1D without modification. In these preclinical tests, DPARC was evaluated under unannounced meals and real-time physiological variability, providing feasibility evidence across simulation, rodent, and swine settings rather than clinical efficacy evidence. These cross-species experiments provide preclinical feasibility evidence for zero-shot transfer across organisms with substantially different glucose-insulin physiology and metabolic dynamics. Finally, we evaluated the physiological alignment of DPARC by showing that its learned latent representations correlated with simulator-derived physiological indicators such as ISF and plasma insulin concentration.

**Table 1.** Overview of experimental stages and key outcomes.

| Stage | Model & deployment | Key methodology | Evaluation metrics | Key findings |
| --- | --- | --- | --- | --- |
| **In silico (n=30)** | UVA/Padova simulator (n=30) | Domain randomization + adaptive RL | TIR, CVGA, COGI, latent-physiology | TIR 72.5% |
| **Swine (n=5)** | Zero-shot deployment in T1D swine (9 sessions) | Policy trained in simulation only | TIR, TAR, TBR under unannounced meals | TIR 86.0%<br>Unannounced standardized meals<br>Exploratory AAPS benchmark<br>No fine-tuning |
| **Rat (n=6)** | Zero-shot deployment in T1D rats (6 sessions) | Policy trained in simulation only | TIR, TAR, TBR under unannounced meals | TIR 79.3%<br>Unannounced standardized meals<br>Rapid-physiology stress test<br>No fine-tuning |
| **Latent analysis** | Simulation data (n=30) + latent decoding | Self-supervised $z \leftrightarrow$ physiology correlation | ISF, CR, BDI regression PCA, sensitivity | Latent $z \leftrightarrow$ ISF ($r=0.93$)<br>$z_3 \leftrightarrow$ plasma insulin ($r=0.59$) |

## RESULTS

### Architecture and operational flow of the DPARC framework

The DPARC leverages an adaptive reinforcement learning framework to enable fully autonomous insulin delivery in T1D. A single policy is trained to generalize across patients in a simulator that introduces variability at two levels: between-patient diversity, achieved by sampling physiological parameters such as insulin sensitivity, carbohydrate absorption kinetics, and endogenous glucose production; and within-patient temporal variability, introduced through time-varying perturbations of these parameters (Fig. 1A). This domain randomization exposes the policy to both routine day-to-day fluctuations and rare extremes. Safety is enforced by design through action caps and hypoglycemia-weighted penalties, producing robust strategies that promote safety. During training, a temporal convolutional encoder (TCN) compresses the preceding 24-h history of CGM, CGM rate of change (CGMdt), and insulin delivery into a latent vector z, which tracks the current metabolic state. This vector conditions an MLP policy network that outputs insulin actions, including basal adjustments and prandial boluses.

**Fig. 1.**
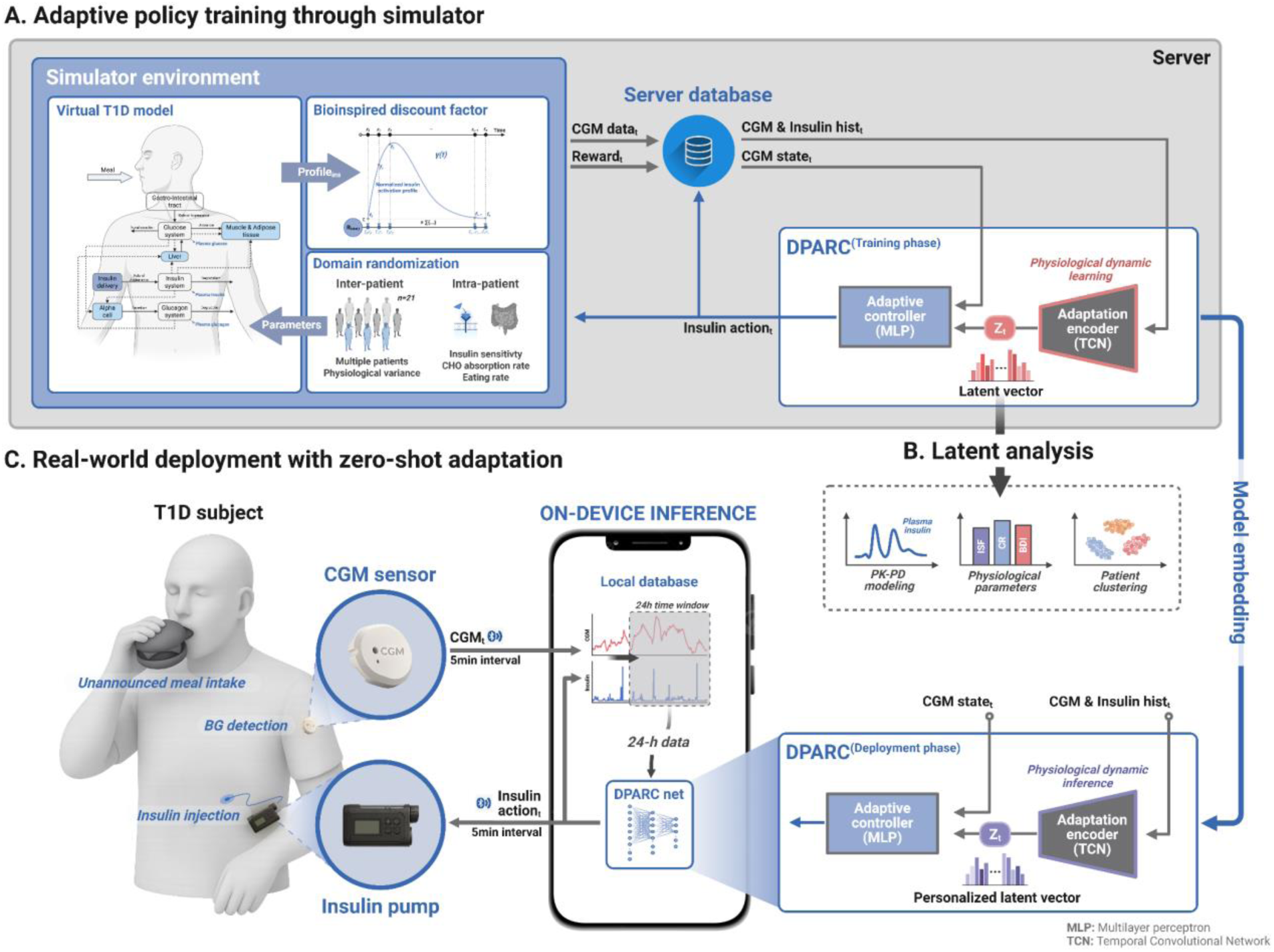
DPARC training, latent-space characterization, and zero-shot deployment. **(A)** Simulator-based training with domain randomization, where inter- and intra-individual variability are introduced into the virtual T1D model (randomized total daily insulin, TDI; insulin sensitivity factor, ISF; time-varying meal absorption kinetics; endogenous glucose production, EGP; and insulin sensitivity, IS). A temporal convolutional network (TCN) encodes the preceding 24-h history of CGM, CGM rate of change (CGMdt), and insulin delivery into a latent physiology vector z, which conditions a multilayer perceptron (MLP) policy network to output insulin actions. A bio-inspired discounting function τ(t) emphasizes near-term safety and stability during training. **(B)** In silico latent-space analysis, where simulation trajectories are projected into z to evaluate alignment with physiological markers (e.g., insulin-sensitivity/absorption proxies), associations with model parameters (e.g., ISF, EGP, IS), and cohort-level organization (individual clustering with within-individual temporal continuity). **(C)** On-device, context-free deployment, where every 5-min cycle updates z from the available CGM and insulin history within a rolling 24-h window, and the policy computes insulin dosing without preset personalization, meal announcements, or user input (zero-shot). Performance is evaluated under unannounced meal conditions, including stochastic in silico meals and supervised preclinical feeding protocols, to assess meal-announcement-free closed-loop operation. During cold start, closed-loop control can begin after 1 h of observed data, with the unobserved portion of the 24-h history window initialized and then progressively replaced by observed CGM and insulin records.

Using simulated trajectories, we analyzed physiological latent vector z over time to characterize the structure and semantics of the latent space (Fig. 1B). The learned representation aligned with physiological markers, with selected latent dimensions correlating with insulin sensitivity and absorption kinetics. It distinguished individuals by physiological profile while preserving within-individual temporal continuity, remaining stable under routine fluctuations yet responsive to acute perturbations. Quantitatively, latent vector z showed strong correlations with physiological proxies, improved clustering separability, and maintained temporal smoothness. These in silico results indicate that latent vector z provides a compact, physiologically meaningful embedding that supports context-free conditioning of the controller while offering post-hoc physiological anchors for subsequent evaluation.

For validation, the trained encoder was fixed and deployed with the policy for real-time control (Fig. 1C). Every 5 min, the system ingests the available CGM and insulin history within a rolling 24-h window, updates latent vector z, and computes dosing decisions without preset personalization parameters, meal announcements, or user input. During cold start, closed-loop operation begins after 1 h of observed data; the remaining unobserved portion of the 24-h window is initialized by the model’s preprocessing procedure and is progressively replaced by real observations. The same model is applied zero-shot across individuals and days, requiring no parameter adjustment. Evaluations under unannounced meals, including stochastic in silico meal schedules and supervised preclinical feeding protocols, demonstrate meal-announcement-free closed-loop operation. A detailed description of the DPARC model architecture and training procedure is provided in Supplementary Note 1, along with schematic diagrams (Supplementary Fig. S1) and implementation details (Supplementary Table S1).

### Performance evaluation of the context-free adaptive policy through in silico experiments

To evaluate the generalization capability of DPARC, we performed in silico validation on an extended cohort of 30 virtual individuals encompassing a wide spectrum of physiological variability, including insulin sensitivity, carbohydrate intake, glucose absorption kinetics, and endogenous glucose production. As comparators, we included TDI-ctx, our previously developed context-conditioned RL controller following Vega et al. (35), which conditions policy outputs on prior knowledge of individual-specific parameters such as TDI. This model represents the most recent RL-based APS validated through a sim-to-real preclinical study and thus provides a strong state-of-the-art benchmark. We also included DPARC-PER (DPARC personalized), a fully personalized upper-bound model trained separately for each individual using the same architecture. In contrast, DPARC operates in a context-free manner, relying solely on latent physiological features inferred in real time by a TCN encoder from the preceding 24 h of CGM data, CGMdt, and insulin-delivery history. This design enabled a direct assessment of DPARC’s ability to adapt across individuals without requiring any predefined user information. Individual daily CGM profiles for each virtual subject are provided in Supplementary Fig. S3.

DPARC maintained greater time in target range (70-180 mg/dL) than TDI-ctx and achieved control comparable to the personalized upper bound, DPARC-PER (Fig. 2A). Cohort-level results (Fig. 2B) confirmed significantly higher TIR and lower time above range (TAR) for DPARC, without an increase in time below range (TBR). Full numerical performance metrics across the cohort are summarized in Supplementary Table S2. In scenarios with unannounced high-carbohydrate meals and time-varying insulin sensitivity, TDI-ctx achieved a mean TIR_70–180_ of 63.4%, while DPARC improved this to 72.5% (*p* < 0.001), approaching DPARC-PER (74.8%). Correspondingly, TAR_180_ decreased from 35.2% (TDI-ctx) to 27.2% (DPARC), compared with 24.7% for DPARC-PER. These results demonstrate that a context-free policy driven by latent physiology can achieve performance close to a personalized upper bound, supporting robust generalization across heterogeneous individual profiles. These findings were consistent across all individual simulations (Supplementary Fig. S3 and Table S2).

**Fig. 2.**
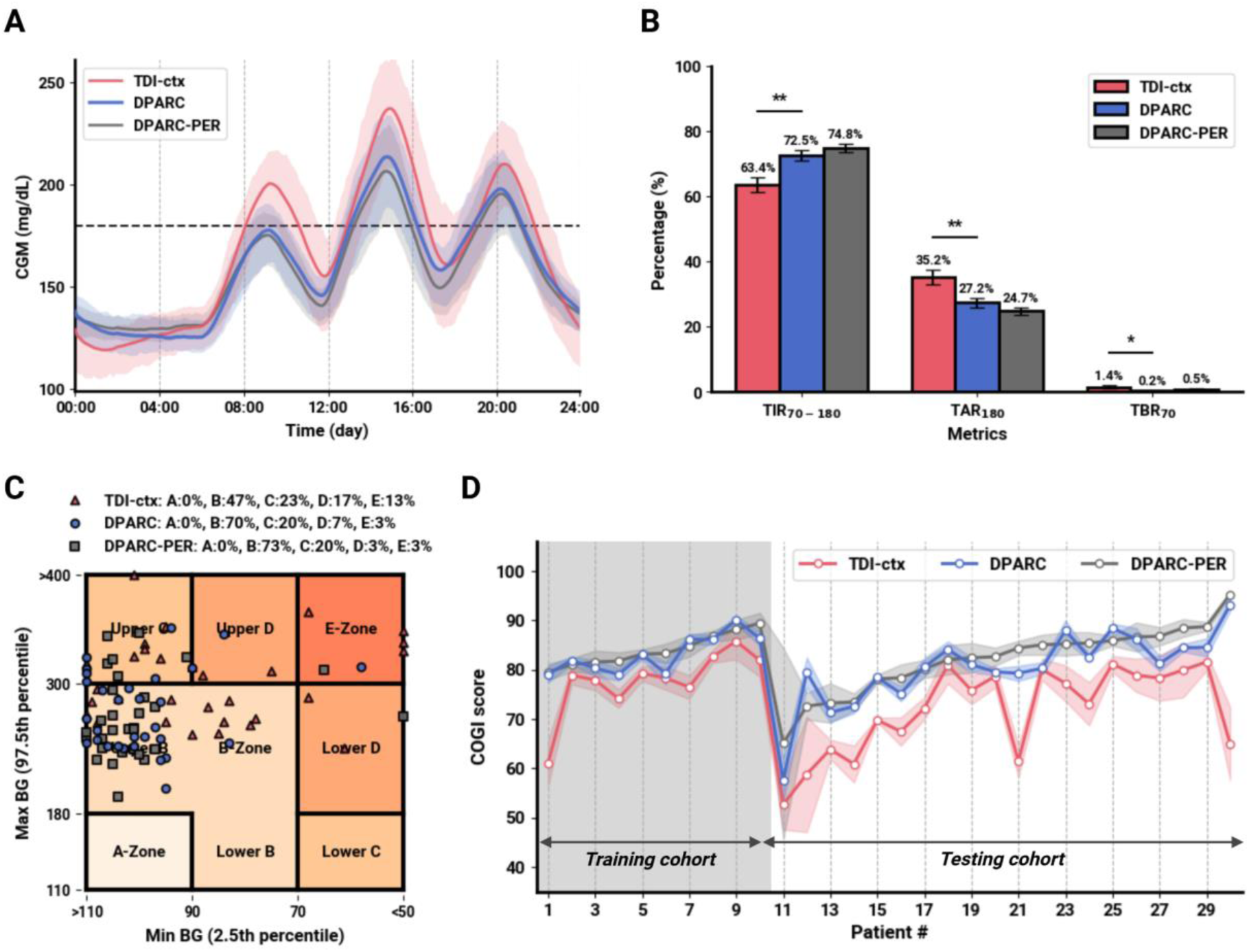
Evaluation of DPARC glucose-control performance in primary in silico generalization experiments under stochastic unannounced meals. **(A)** Mean±SD CGM profiles across 30 virtual individuals comparing the proposed DPARC model, the baseline TDI-conditioned model (TDI-ctx), and the fully personalized upper-bound model (DPARC-PER). Meal timing, carbohydrate amount, and meal absorption were randomized, and each meal had a 10% probability of being skipped; no meal information was provided to the controller. **(B)** DPARC significantly improved glycemic metrics, including time in range (TIR), time above range (TAR), and time below range (TBR), relative to TDI-ctx, achieving performance close to the personalized DPARC-PER. **(C)** Control Variability Grid Analysis (CVGA) showed that DPARC increased the proportion of glucose readings within the normoglycemic B-zone while reducing entries in clinically dangerous zones. **(D)** Individual-level evaluation using Composite Continuous Glucose Monitoring Index (COGI) scores demonstrated that DPARC maintained consistent performance across both training and test cohorts, supporting its preclinical feasibility as a single context-free policy. Error bars and shaded regions represent mean±SD.

In terms of safety, DPARC also showed marked improvement. TBR70 decreased from 1.4% (TDI-ctx) to 0.2% (DPARC), and Level 2 hypoglycemia exposure, defined as CGM <54 mg/dL, was eliminated in this simulation setting (TBR_54_ = 0.0%), matching DPARC-PER. These outcomes suggest that a context-free adaptive policy can approach the safety profile of fully personalized models in simulation without prior individual configuration.

The advantage of DPARC was further supported by Control Variability Grid Analysis (CVGA) (*36*) (Fig. 2C), which simultaneously evaluates the safety and stability of the proposed controller. DPARC placed 70% of points in the B-zone (normoglycemia) versus 47% for TDI-ctx, while reducing dangerous zones (D+E) from 30% to 10% (*p* < 0.001), demonstrating significant improvement in safety-critical outcomes. Importantly, DPARC maintained consistent performance across individuals (Fig. 2D). Individual-level Composite Continuous Glucose Monitoring Index (COGI) scores (*37*) showed that DPARC delivered stable control across the cohort, whereas DPARC-PER, despite achieving optimal performance for some individuals, exhibited greater variability. This suggests that DPARC not only generalizes across diverse physiology but also ensures reliability at the individual level—a crucial advantage for scalable deployment in clinical practice.

### Cold-start deployment in silico

Having established cohort-level generalization, we next evaluated DPARC’s cold-start robustness at deployment by initializing with minimal context and assessing its stabilization dynamics. For each of 30 virtual individuals, we collected 1 h of open-loop history before switching to fully closed-loop control without meal announcements or user input. The same policy was applied zero-shot across all individuals, with no per-individual tuning. Control executed at 5-min intervals and operated with a rolling 24-h window that progressively filled during day 1; at switchover, only 1 h of history was available. Across three consecutive days, control remained stable without oscillatory drift or instability. Aggregate traces showed rapid correction of postprandial excursions and a progressive reduction in variability over time, with CGM standard deviation decreasing across days and nocturnal periods becoming steadier (Fig. 3A).

**Fig. 3.**
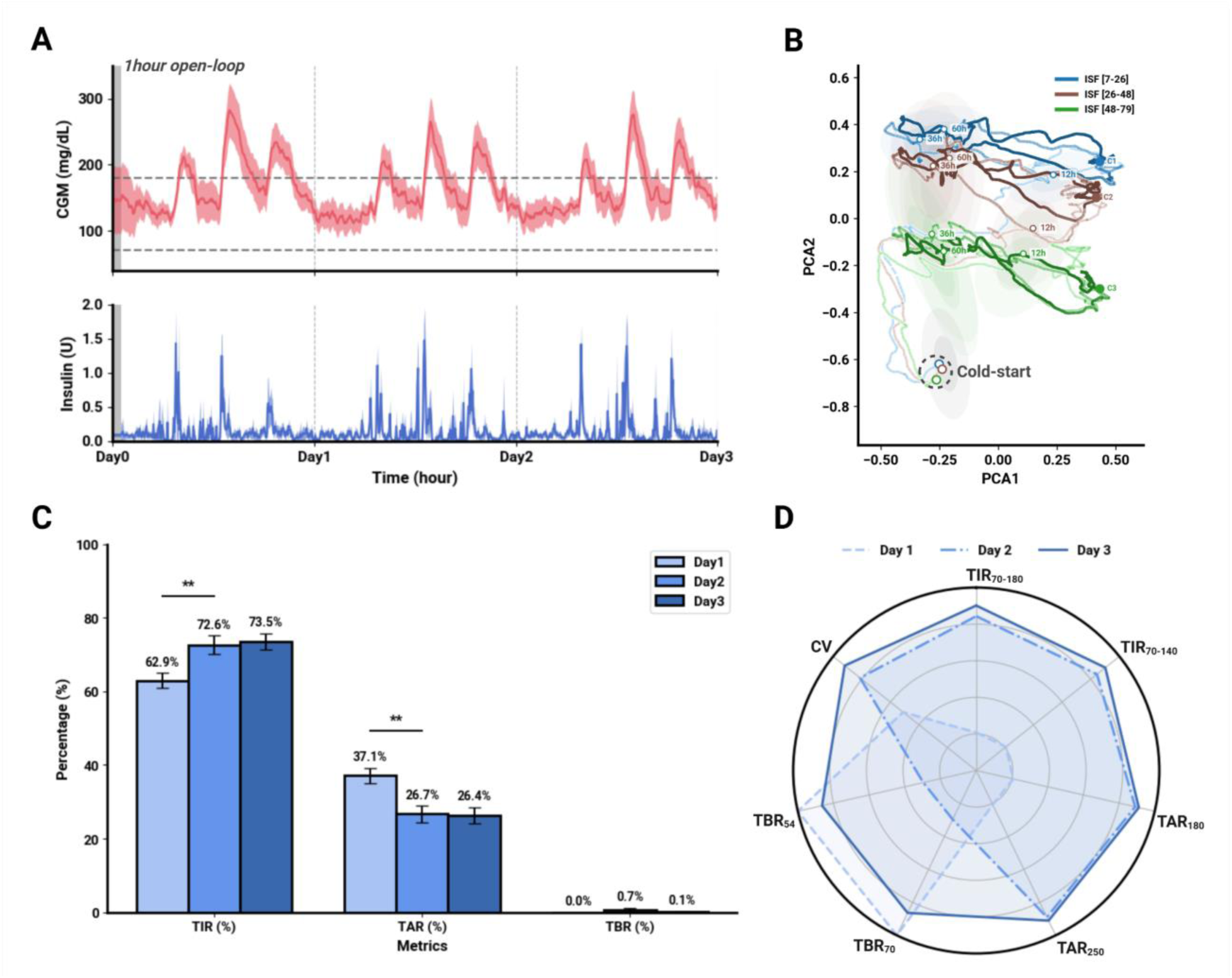
Cold-start deployment and day-over-day stabilization. **(A)** CGM and insulin Mean±SD traces for 30 virtual individuals over 3 days. Control begins after a 1-h open-loop period, with variability decreasing over time and postprandial excursions reduced by day 3. **(B)** Latent-space trajectories grouped by insulin-sensitivity bands. Individuals start within a compact cold-start region, form separable clusters by approximately 12 h, and subsequently evolve within cluster-specific neighborhoods. **(C)** Day-wise outcomes showing increasing TIR from days 1 to 3, decreasing TAR, and consistently low TBR, with a transient rise on day 2 that resolves by day 3. Error bars represent inter-individual variability. **(D)** Radar plot summarizing multiple metrics (TIR_70–180_, TIR_70–140_, TAR_180_, TAR_250_, TBR_70_, TBR_54_, and coefficient of variation). The shaded area expands and becomes more uniform from days 1 to 3, indicating broad stabilization without compromising safety.

To examine internal adaptation, we analyzed encoder latent trajectories. Individuals were grouped into three bands by insulin sensitivity factor (ISF 7–26, 26–48, and 48–79 (mg/dL/U)). Immediately after the cold start, trajectories originated from a compact low-variance region reflecting limited context. By approximately 12 h, they separated into clusters corresponding to sensitivity bands and subsequently evolved smoothly within cluster neighborhoods while maintaining within-individual continuity (Fig. 3B). Within-cluster drift aligned with meal timing and circadian segments, suggesting that the encoder rapidly identifies an individualized operating point and then updates it smoothly as physiology evolves. For visualization, a two-dimensional projection capturing the majority of variance was used, making both early convergence and subsequent local motion apparent.

Performance metrics improved in parallel with latent organization. Mean TIR increased from 62.9% on day 1 to 72.6% on day 2 and 73.5% on day 3, while TAR decreased from 37.1% to 26.7% and 26.4%. TBR remained low overall, with a transient rise on day 2 that resolved by day 3 (0.0%, 0.7%, and 0.1%; Fig. 3C). Narrow-band TIR and coefficient of variation followed similar trends, indicating tighter control rather than a trade-off across endpoints. A radar summary confirmed simultaneous gains across multiple metrics and day-over-day stabilization (Fig. 3D), demonstrating DPARC’s continual adaptation capability as it progressively refines physiological understanding through real-time interaction. Collectively, these findings demonstrate that DPARC stabilizes from a cold start using only limited history, converges to individualized latent vector spaces within half a day, and sustains stable in silico control thereafter, supporting the preclinical feasibility of rapid startup with minimal initial data. Additional robustness testing under perturbed basal infusion rates (80%, 100%, and 120% of nominal) further confirmed stability of control during cold-start conditions (Supplementary Fig. S4).

### Adaptive response performance to real-time changes in dynamic physiology

Current automated insulin delivery (AID) systems that rely on fixed personalization parameters exhibit inherent limitations in handling rapid and unpredictable physiological changes. To evaluate the capacity of DPARC for real-time adaptation under such challenging conditions, we designed a progressive perturbation experiment in which insulin sensitivity (IS) was gradually varied within a single virtual individual. Specifically, IS was gradually shifted from -50% to +50% over 8 days under a controlled fixed-meal protocol without meal announcements (Fig. 4A). This scenario reflects clinical contexts such as infection, stress, or hormonal fluctuation, introducing extreme yet clinically plausible variability. Despite these substantial perturbations, DPARC maintained stable mean BG and TDI through real-time compensatory adjustments (Fig. 4B). By contrast, the context-based TDI-ctx model failed to adapt, resulting in progressive hyperglycemia and declining performance as IS changed. These results demonstrate DPARC’s ability to dynamically adjust to latent physiological changes in real-world-like conditions, overcoming a key limitation of conventional fixed-parameter controllers. To further examine the internal adaptation mechanism of DPARC, we visualized the temporal evolution of latent vector z using two-dimensional principal component analysis (PCA) (Fig. 4C). The resulting trajectories revealed structured and continuous transitions in latent space aligned with shifting IS, providing direct evidence that the TCN encoder captures and adapts to unobservable physiological dynamics in real time.

**Fig. 4.**
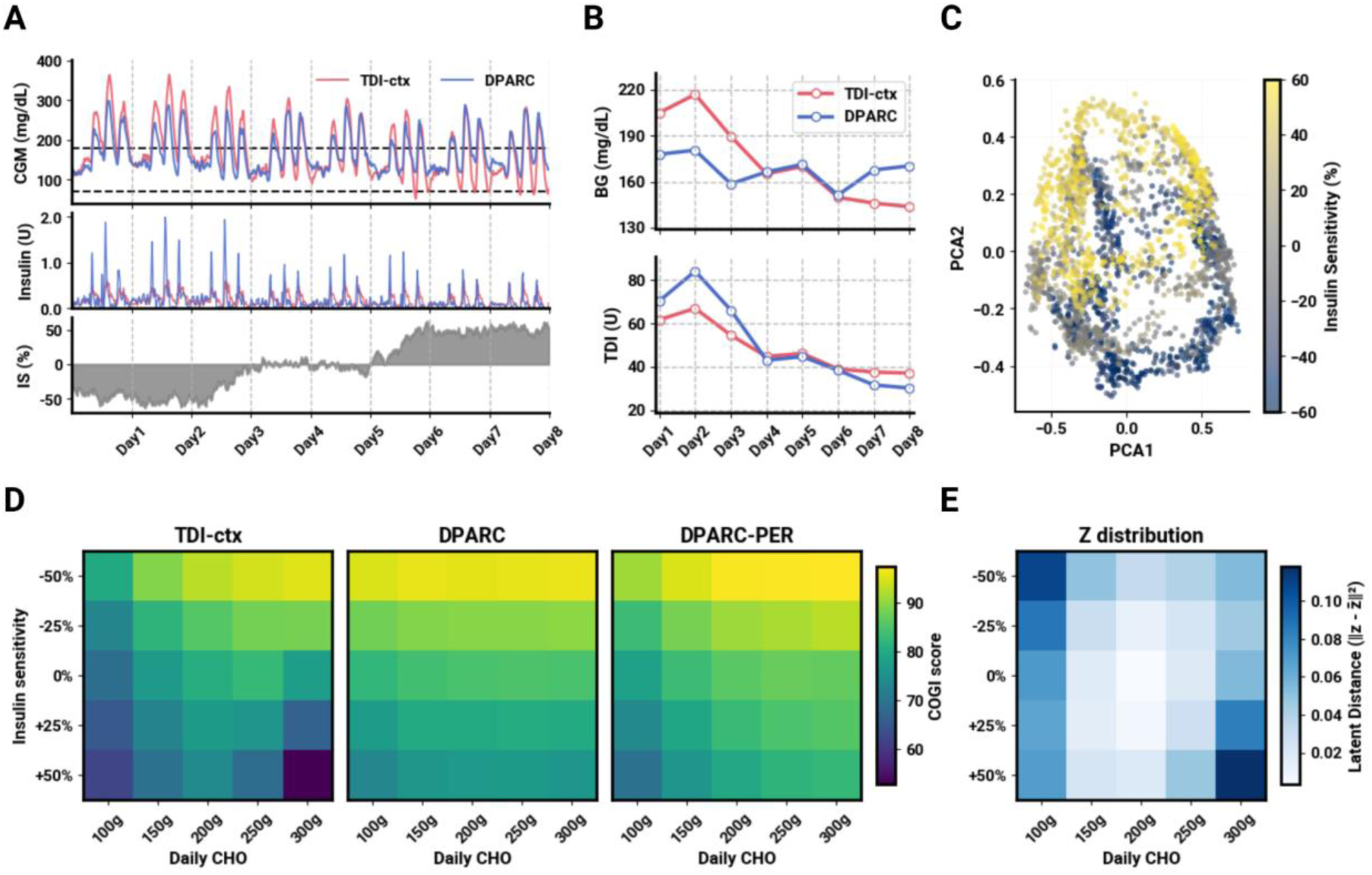
Evaluation of DPARC’s real-time adaptive capability to physiological variability. **(A)** In a progressive perturbation scenario where insulin sensitivity (IS) shifts linearly from – 50% to +50% over 8 days, DPARC maintains stable glucose control and dynamically adjusts insulin dosing in real time. **(B)** Despite changing IS, DPARC sustains stable mean blood glucose (BG) and TDI, while TDI-ctx fails to adapt, leading to progressive hyperglycemia**. (C)** Principal component analysis (PCA) visualization of latent representations extracted by DPARC shows smooth trajectory shifts in latent space corresponding to IS variation, indicating effective tracking of internal physiological changes. **(D)** Across the full combinatorial grid of IS and carbohydrate intake (CHO) in three virtual individuals, DPARC consistently achieves higher COGI scores than both TDI-ctx and the personalized DPARC-PER model. **(E)** Latent-distance analysis confirms smooth, structured adjustments in representation space in response to varying physiological parameters.

For a more comprehensive assessment of generalization, we selected three virtual individuals spanning a wide range of total daily insulin requirements and evaluated DPARC across the full grid of insulin sensitivity and carbohydrate intake perturbations. Performance was quantified using the COGI score (Fig. 4D). Across this broad physiological space, DPARC consistently maintained high performance with only shallow degradation, outperforming TDI-ctx throughout the grid and especially under low-sensitivity and high-carbohydrate conditions where TDI-ctx control deteriorated substantially. Notably, DPARC also matched or exceeded the personalized upper bound (DPARC-PER) in several extreme regions, underscoring its superior adaptability under physiologically challenging conditions. To investigate the basis of this robustness, we visualized pairwise Euclidean distances between latent features extracted under each IS–CHO condition, generating a latent distance heatmap (Fig. 4E). The resulting structure exhibited gradual and systematic movement patterns in latent space as IS and CHO increased, indicating that the encoder’s internal representations adapt in a smooth and physiologically meaningful manner. Importantly, the tight correspondence between COGI scores and latent-space transitions suggests that the encoder reliably captures core physiological variables such as insulin sensitivity, carbohydrate absorption rate, and endogenous glucose production. These latent representations are subsequently passed to the adaptive controller, enabling quantitatively aligned insulin-delivery decisions. The emergence of a self-organizing latent structure, achieved without external supervision or predefined parameters, provides the foundation for DPARC’s real-time physiological alignment and adaptive control capabilities. In addition, we tested DPARC under acute, real-world–relevant perturbations such as exercise-induced increases in insulin sensitivity, post-prandial exercise, and post-prandial insulin resistance. In all scenarios, DPARC promptly stabilized glucose without loss of safety (Supplementary Fig. S5).

### Self-supervised latent encoding reveals physiologically aligned latent dynamics

The core technological distinction of DPARC lies in its ability to learn latent representations that infer an individual’s physiology in real time without relying on externally predefined personalization parameters. Using in silico trajectories (Fig. 5), we analyzed the latent vectors produced by the TCN encoder and their influence on policy decisions. The encoder processes CGM and insulin-infusion histories from the preceding 24-h and updates a latent physiology vector every 5 min via a temporal convolutional network (Fig. 5A). This latent vector simultaneously encodes relatively static physiological traits that vary slowly (for example, ISF and carbohydrate ratio, CR) and dynamic physiology that changes rapidly (for example, plasma insulin proxies), forming the substrate for context-free adaptive control. To quantify how each latent dimension affects the controller, we conducted a counterfactual perturbation analysis in which small, fixed step changes were applied to individual latent dimensions, and the resulting shifts in insulin actions were measured across time and across individuals. Latent dimensions z_3_, z_8_, and z_27_ exerted statistically reliable influence on dosing (Fig. 5B), with consistent effects across a range of perturbation magnitudes. This finding indicates that the latent space encodes variables directly conditioning control rather than serving merely as a compressed representation. Qualitatively, perturbations along these dimensions altered the controller’s aggressiveness following meals and its conservativeness in low-glucose scenarios, demonstrating a direct pathway from physiological state estimation to safety-aware action selection.

**Fig. 5.**
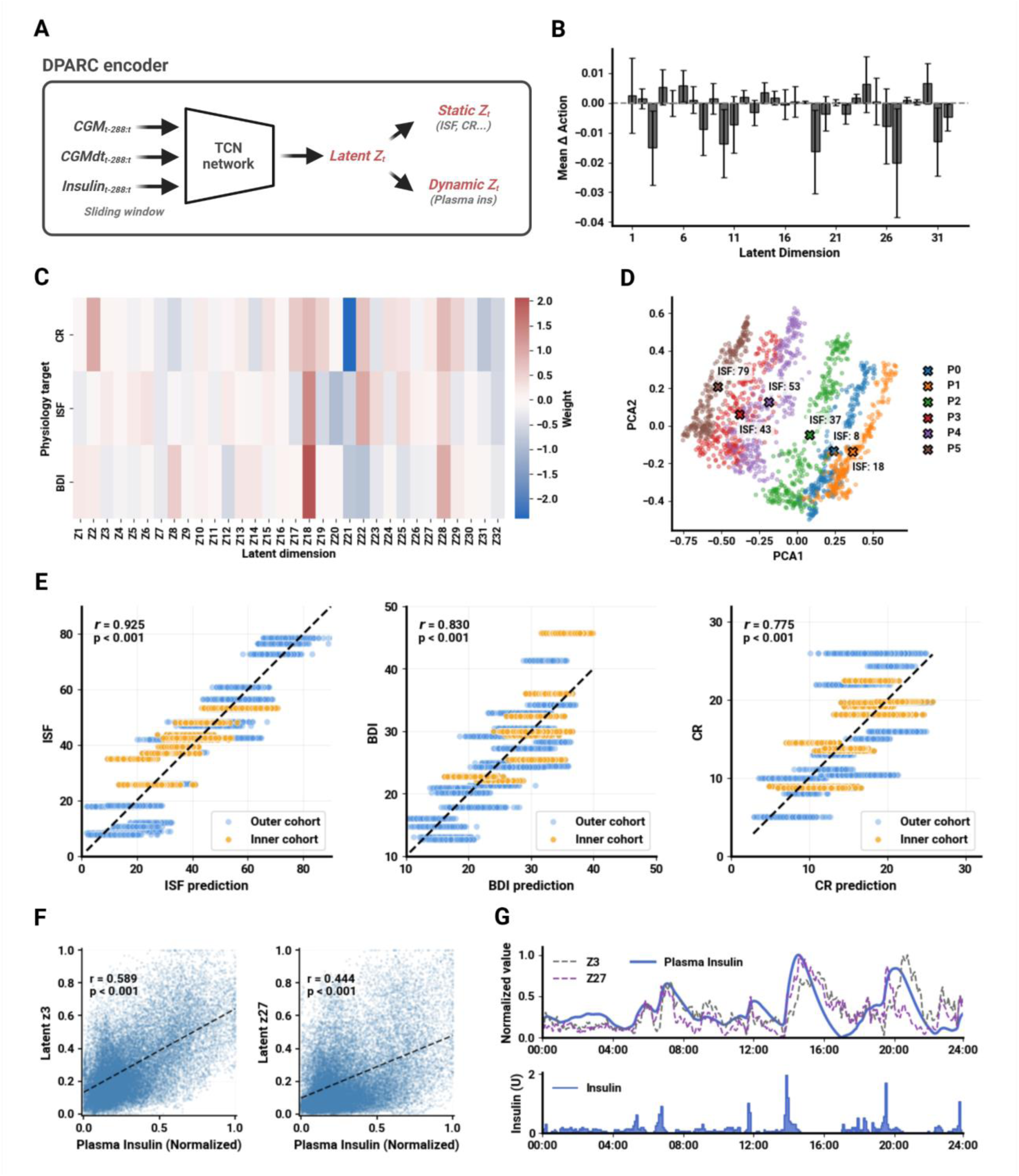
Physiological alignment of latent representations learned by the DPARC encoder. **(A)** Encoder architecture. A 24-h sliding window of CGM, CGMdt, and insulin-infusion history is processed by a TCN to generate a latent vector z. This embedding captures long-term physiological traits such as ISF and carbohydrate ratio (CR), as well as short-term metabolic dynamics such as plasma insulin kinetics. **(B)** Counterfactual sensitivity analysis of policy behavior. Small perturbations applied to individual latent dimensions revealed that several, including z_3_, z_8_, and z_27_, exert significant influence on insulin-dosing decisions. **(C)** Regression analysis between latent vectors and physiological targets (ISF, basal daily insulin [BDI], and CR) showed that each target parameter is distributed across multiple latent dimensions, indicating compositional rather than one-to-one encoding. **(D)** Principal component projection of latent vectors from representative virtual individuals with distinct ISF values produced well-separated clusters, demonstrating structural differentiation of intrinsic physiological profiles. **(E)** Correlations between physiology inferred from latent vectors and simulator ground truth were high (ISF: r = 0.925; BDI: r = 0.830; CR: r = 0.775), showing that these simulator-derived parameters are predictable from the learned embedding without explicit supervision. **(F)** Latent dimensions z_3_ and z_27_ correlated significantly with plasma insulin concentrations (r = 0.589 and r = 0.444, respectively; both *p* < 0.001), suggesting that typically unobservable metabolic indicators can be inferred indirectly. **(G)** In silico simulations further showed that the temporal trajectories of z_3_ and z_27_ closely tracked plasma insulin curves, indicating that the encoder captures pharmacokinetic dynamics over time.

Subsequently, we examined how known physiological parameters map onto the latent space. Using linear probe models, ISF, basal daily insulin (BDI), and CR exhibited distributed many-to-many correspondence across multiple latent dimensions (Fig. 5C), rather than collapsing onto single coordinates. A projection of latent vectors from five representative individuals produced clearly separable clusters while preserving within-individual temporal continuity (Fig. 5D), consistent with an encoder that captures both stable inter-individual physiology and dynamic intra-individual trajectories. Probe predictions were highly correlated with simulator ground truth—ISF: r = 0.925, BDI: r = 0.830, and CR: r = 0.775, all *p* < 0.001 (Fig. 5E)—demonstrating that physiological parameters are linearly predictable from the learned embedding despite the absence of parameter labels during training. For dynamic physiology, latent dimensions z_3_ and z_27_ tracked simulated plasma-insulin time courses with correlations of 0.589 and 0.444, respectively (both *p* < 0.001; Fig. 5F), and their temporal evolution aligned with diurnal fluctuations in meals and insulin dosing in silico (Fig. 5G). Collectively, these results indicate that DPARC learns a self-organizing, physiologically aligned latent space that captures slow-varying parameters and fast PK-like dynamics and links these latent features to dosing actions. These analyses provide post-hoc explanatory anchors, but they do not make the neural policy fully mechanistically interpretable.

### Rodent rapid-physiology stress test of zero-shot DPARC deployment

Before validating DPARC in a clinically relevant large animal model, we first challenged the DPARC’s adaptability in a rodent model (n=6, streptozotocin-induced diabetic Sprague-Dawley rats). Rodents possess significantly faster glucose turnover rates and insulin kinetics compared to humans or swine, presenting a harsher control environment characterized by rapid glycemic fluctuations. Validating the algorithm in this high-frequency domain was crucial to verify whether the TCN encoder could capture and react to fast-evolving physiological states in a zero-shot manner.

The simulation-trained DPARC policy was deployed directly to the rats following the zero-shot protocol outlined in Fig. 6A, without any species-specific tuning or open-loop warm-up periods. Despite the substantial mismatch between the in silico environment and the in vivo deployment setting (Fig. 6B), DPARC executed closed-loop control without species-specific tuning. As shown in the representative 24-h glucose and insulin trajectories (Fig. 6C), DPARC responded to postprandial excursions under unannounced meals and rapid metabolic conditions. Quantitatively, DPARC achieved a mean TIR of 79.3%, with TAR of 15.2% and TBR_70_ of 5.6% (Fig. 6D). Individual subject traces and detailed performance metrics are provided in Supplementary Fig. S9 and Table S3. These results are interpreted as a rapid-physiology stress test rather than as evidence of clinically acceptable glycemic safety. The observed TBR70 of 5.6% indicates that additional safety tuning is required before human translation.

**Fig. 6.**
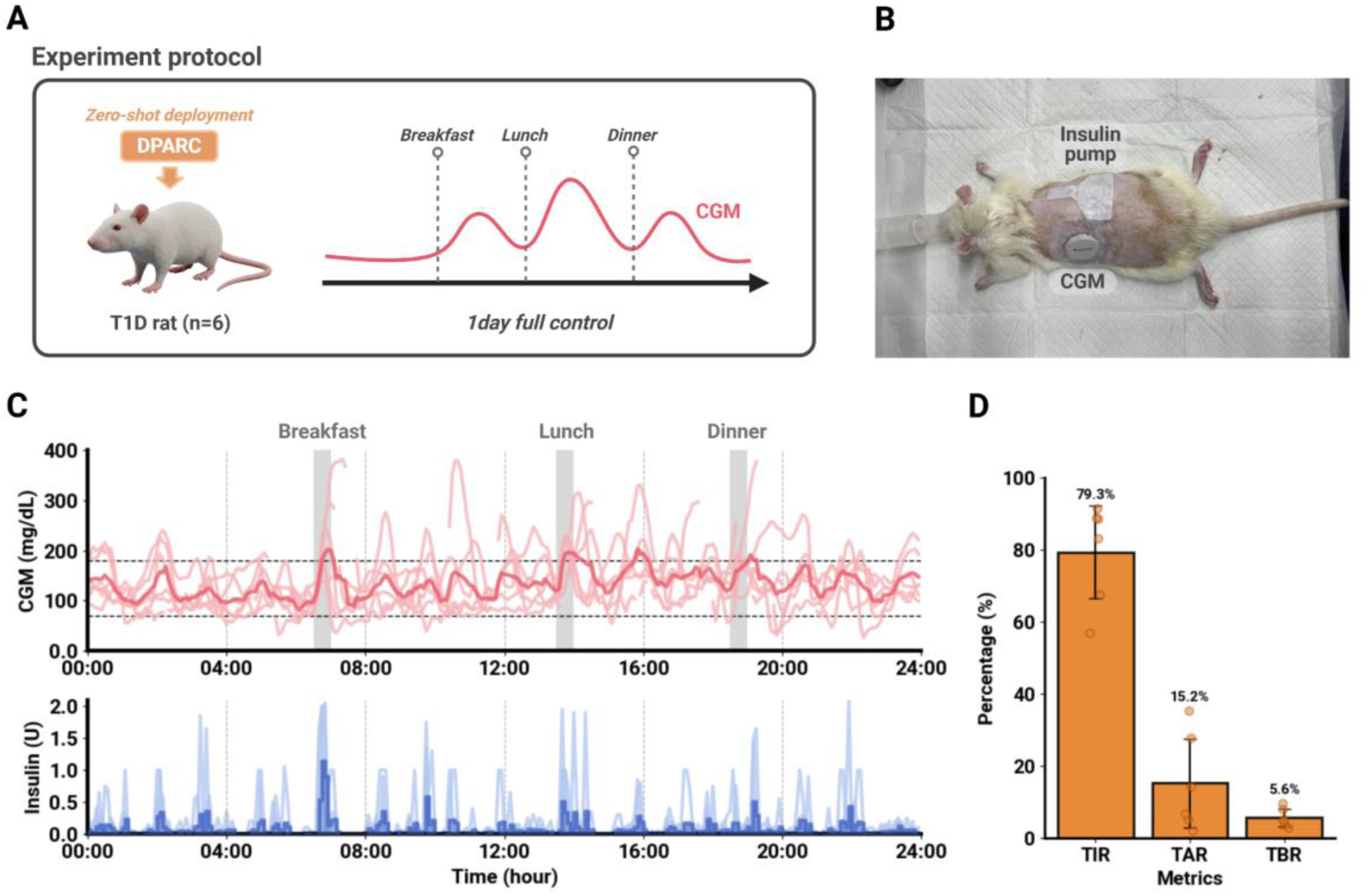
Rodent rapid-physiology stress test of zero-shot DPARC deployment. **(A)** Protocol. A policy trained entirely in simulation was deployed without modification (zero-shot) in six streptozotocin-induced diabetic Sprague-Dawley rats (n=6). Fully closed-loop control was executed for 24 h under a standardized three-meal protocol without meal announcements at a 5-min sampling interval to test deployment under rapid metabolic fluctuations. **(B)** Experimental setting. Rats were instrumented with a subcutaneous insulin catheter and a Dexcom G7 continuous glucose monitor, both positioned dorsally to ensure stability during freely moving conditions. **(C)** 24-h control. Top: CGM glucose traces from individual sessions showing handling of rapid dynamics. Bottom: corresponding insulin infusion profiles. DPARC responded to postprandial excursions under the rapid physiological kinetics of the rodent model. **(D)** Quantitative outcomes. DPARC achieved a mean TIR of 79.3%, TAR was 15.2%, and TBR_70_ was 5.6%, indicating that the rodent study should be interpreted as a stress test rather than as clinical safety evidence. Bars represent group means; individual data points shown with error bars representing SD.

### Zero-shot sim-to-real deployment in swine with exploratory benchmark comparison

Bridging simulation and in vivo physiology remains a central challenge for AI-based medical systems. To evaluate the proposed DPARC’s zero-shot operation without manual tuning, we conducted preclinical studies in Yorkshire swine with chemically induced T1D (n = 5; 20–40 kg). The trained DPARC algorithm was deployed directly, without modification, parameter calibration, or prior personalization (Fig. 7A). Animals were maintained on a daily schedule of breakfast, lunch, and dinner, but meal information was not provided to the algorithm, and no manual boluses were administered. Thus, DPARC functioned in a fully closed-loop manner without any user input (Fig. 7B). Real-time control was executed at 5-min intervals using CGM and insulin-infusion histories, with built-in safety constraints active throughout all sessions to prevent overdosing.

**Fig. 7.**
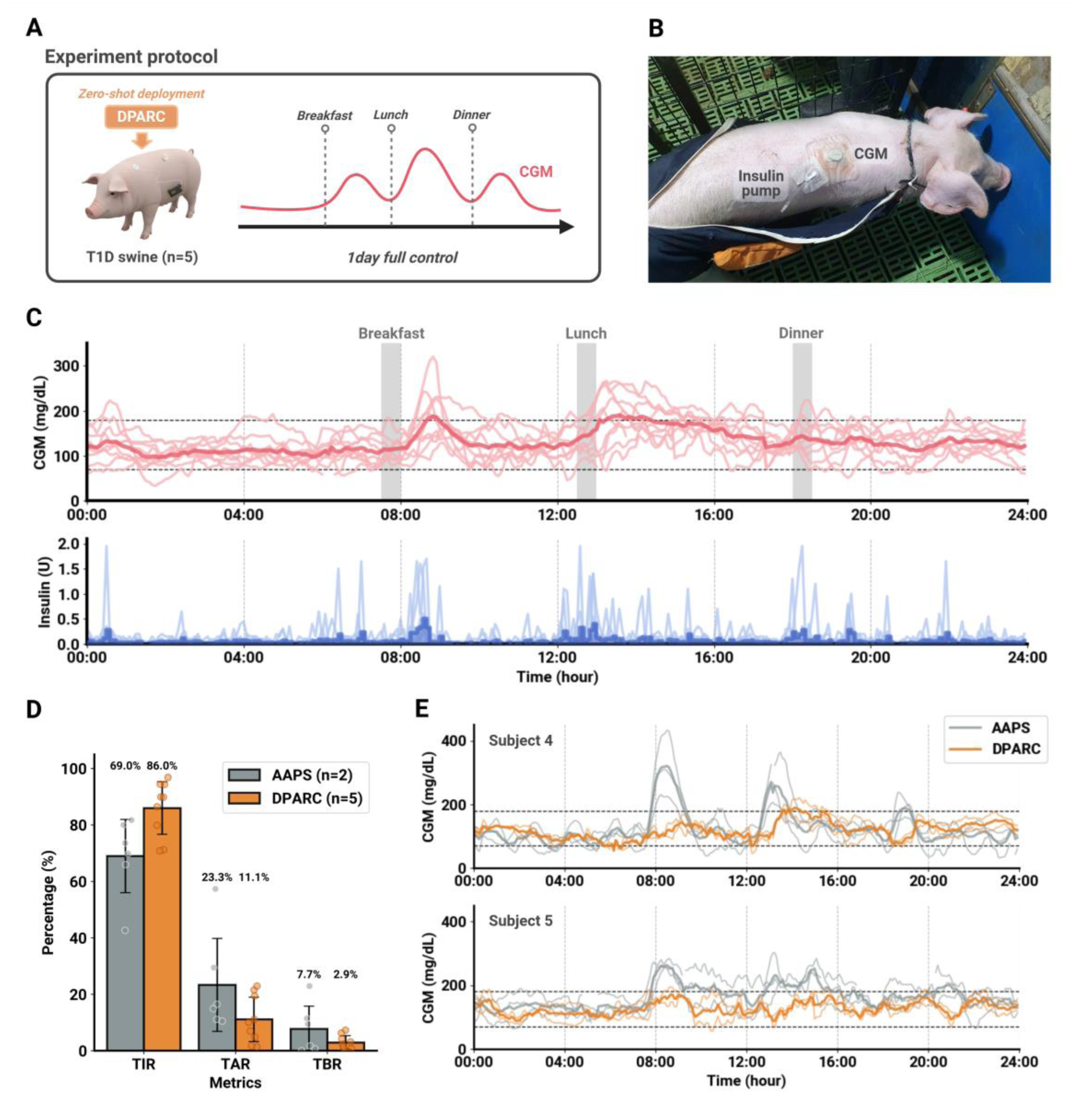
Zero-shot sim-to-real deployment in Yorkshire swine with exploratory AAPS benchmark comparison. **(A)** Protocol. A policy trained entirely in simulation was deployed without modification (zero-shot) in five Yorkshire swine with chemically induced T1D. Fully closed-loop control was executed under a three-meal schedule (no meal announcements) at a 5-min sampling interval. A subset of two animals (Subjects 4 and 5) underwent crossover testing comparing AAPS and DPARC. **(B)** Experimental setting. Swine were instrumented with a subcutaneous insulin catheter and a Dexcom G7 continuous glucose monitor, both positioned dorsally to ensure stability during freely moving conditions. **(C)** 24-hour control. Top: CGM glucose traces from individual sessions with cohort mean and variability band. Bottom: corresponding insulin infusion profiles. DPARC executed closed-loop regulation and responded to postprandial excursions. **(D)** Comparative performance metrics. DPARC feasibility results are shown across five animals and nine sessions, whereas the AAPS comparison was restricted to an exploratory crossover subset of two animals and six AAPS sessions. In this subset, mean TIR was higher under DPARC than AAPS: 86.0% vs. 69.0%. TAR decreased from 23.3% (AAPS) to 11.1% (DPARC), while TBR remained low for both systems (2.9% vs. 7.7%, not significant). Individual data points shown; error bars represent SD. **(E)** Exploratory crossover traces. Within-subject CGM traces for Subjects 4 and 5 under AAPS (gray) and DPARC (orange) illustrate qualitative differences in glucose regulation under identical meal conditions.

Across multiple 24-h sessions, DPARC demonstrated feasibility of zero-shot sim-to-real deployment in swine. Glucose trajectories showed closed-loop regulation across sessions, with insulin delivery responding to postprandial excursions (Fig. 7C). Detailed performance metrics for each animal and session are summarized in Supplementary Table S4. Level 2 hypoglycemia exposure, defined as CGM <54 mg/dL, averaged 0.5% across swine sessions. We therefore report this as a low but nonzero hypoglycemia burden rather than as negligible. Furthermore, session-to-session variability was low, reflecting stable and reproducible performance across repeated deployments. Without any predefined personalization parameters or user calibration, DPARC adapted autonomously from CGM and insulin-infusion histories, using a single zero-shot policy across all animals without individualization. Individual glucose and insulin traces for each swine are provided in Supplementary Fig. S11.

To benchmark DPARC against established automated insulin delivery systems, we performed a direct comparison with the Android Artificial Pancreas System (AAPS), an open-source algorithm widely validated in preclinical and clinical settings (42). In this study, AAPS was configured for FCL operation by enabling its advanced Super Micro Bolus (SMB) functionality, allowing it to deliver automated correction boluses without user intervention. AAPS relies on user-specific profile settings such as basal rate, ISF, and carbohydrate ratio, whereas DPARC infers latent physiological context from CGM and insulin history without receiving these profile parameters. Although the SMB algorithm can deliver automated corrections, its dosing magnitude is mathematically bound by the pre-programmed ISF and carbohydrate ratio.

Consequently, if these fixed settings deviate from the subject’s real-time physiological state, as often occurs during unannounced meals or stress, the controller cannot adapt its response intensity. This difference allowed us to compare a profile-based open-source AID approach with a parameter-free physiology-history-conditioned policy under the same preclinical protocol.

In an exploratory crossover subset involving Subjects 4 and 5, DPARC showed higher mean TIR than AAPS under the tested preclinical protocol (86.0% vs. 69.0%, Fig. 7D), with lower TAR180 (11.1% vs. 23.3%). TBR70 was also lower under DPARC than AAPS in this subset (2.9% vs. 7.7%). Because the AAPS comparator arm included only two animals and multiple sessions per animal, this comparison is presented as a hypothesis-generating large-animal benchmark rather than definitive evidence of statistical superiority. Within-subject traces from the crossover subset are shown to illustrate qualitative differences in glucose regulation under identical meal conditions (Fig. 7E).

## DISCUSSION

Our results demonstrate that reinforcement learning with adaptive latent representations can overcome fundamental limitations that have constrained the deployment of truly autonomous APS for over two decades. The most advanced systems in automated insulin delivery among currently available commercial options, including Medtronic MiniMed 780G and Tandem Control-IQ, require manual configuration of personalization parameters such as TDI, CR, and ISF, with studies showing that parameter estimation errors can compromise glycemic control and increase glucose variability (*38*). In contrast, the proposed DPARC achieved stable glycemic control without any predefined parameters, instead inferring physiological state directly from 24-h CGM and insulin histories. This parameter-free approach addresses a critical clinical need, as hybrid closed-loop systems require subjects to count carbohydrates, a process that is both burdensome and error-prone (*39*). By eliminating these requirements, our approach may substantially reduce the clinical burden associated with APS deployment and maintenance, particularly for vulnerable populations including pediatric people with diabetes and older adults who face greater challenges with device management.

DPARC’s parameter-free design is enabled by three innovations that distinguish it from existing approaches. The pharmacokinetics-informed reward structure emphasizes long-horizon objectives that account for insulin action delays rather than immediate corrections, aligning policy optimization with clinically meaningful stability outcomes. Domain randomization during training exposes the policy to inter- and intra-individual variability in insulin sensitivity, carbohydrate absorption kinetics, endogenous glucose production, and other physiological factors, improving robustness to distributional shifts during deployment that have limited other simulation-trained controllers. Most critically, the self-supervised latent representation enables physiology-aware control without requiring labeled clinical parameters, providing personalization benefits without explicit parameter entry or recalibration.

The physiological alignment of DPARC’s latent representations provides post-hoc explanatory anchors for the learned controller. Our in silico analyses showed that selected latent dimensions correlate with simulator-derived parameters such as ISF, BDI, and CR, while other dimensions track dynamic postprandial or pharmacokinetic-like physiology. These findings suggest that the encoder learns physiologically meaningful structure rather than an arbitrary compressed state. However, these correlations do not make the full neural policy mechanistically interpretable; instead, they provide partial explanatory anchors that can support subsequent safety analysis and hypothesis generation.

The rodent experiment was designed as a rapid-physiology stress test of zero-shot deployment rather than as a demonstration of clinically acceptable glycemic safety. Rodent glucose-insulin dynamics are faster and more volatile than human physiology, creating a challenging closed-loop setting. In this context, DPARC achieved a mean TIR of 79.3% with TAR of 15.2%, but TBR70 reached 5.6%. We therefore interpret the rat data as evidence that the frozen policy can operate under a highly dynamic in vivo physiology, while also highlighting the need for additional safety tuning and supervised evaluation before human translation.

The swine experiments provided a large-animal feasibility test of zero-shot DPARC deployment using the same frozen policy without subject-specific tuning. In the exploratory two-animal crossover subset, DPARC showed higher mean TIR and lower TAR than AAPS under the tested protocol. However, because the comparator arm included only two animals and repeated sessions, these data should be interpreted as hypothesis-generating rather than definitive evidence of superiority over AAPS. The value of this experiment is that it demonstrates feasibility of parameter-free, meal-announcement-free control in a large-animal model and motivates more rigorous prospective benchmarking, including larger animal cohorts and supervised human studies.

Beyond glycemic metrics, the parameter-free design of DPARC fundamentally redefines the accessibility of automated diabetes care. Studies highlight that user burden remains a primary barrier to APS adoption (46), particularly for pediatric users, older adults, and individuals with cognitive or functional limitations. High-performance systems that require complex setup and extensive knowledge can inadvertently widen the digital divide in healthcare. Current systems require extensive initial setup and ongoing recalibration, reducing uptake and exacerbating disparities in technology adoption. By decoupling optimal control from the need for expert configuration, DPARC minimizes the cognitive load on patients and caregivers, effectively lowering the barrier to entry. This approach may expand access to fully automated systems for populations historically underserved by diabetes technologies. When considered alongside our in silico, multi-species preclinical, and latent physiological-alignment analyses, these findings support DPARC as a preclinical proof-of-concept framework and provide a rationale for supervised prospective human evaluation. Because our in vivo studies remain limited in scale and do not include prospective human deployment, the results should be interpreted as preclinical feasibility evidence rather than clinical efficacy evidence. Building on this foundation, we are preparing supervised clinical evaluation under appropriate regulatory and ethical oversight; clinical efficacy and readiness for unsupervised human use remain to be established in future studies.

## MATERIALS AND METHODS

### DPARC algorithm architecture: A paradigm shift in personalization

A fundamental limitation of existing AID systems is their intrinsic dependence on manually preset individual-specific parameters. All current commercial and investigational systems require extended initialization periods, personalized basal insulin settings, and frequent manual tuning. This reliance not only conflicts with the vision of fully autonomous glucose control but also limits clinical practicality.

The proposed algorithm, DPARC, addresses these constraints by introducing a paradigm of *physiology-aware latent representation learning*. Our central insight is that rather than explicitly modeling each user’s physiological parameters, dynamic physiological features can be inferred directly from TCN-processed glucose-insulin trajectories. This strategy overcomes the limitations of traditional model-based controllers and enables real-time adaptation to intra-individual physiological variability.

The temporal encoder of DPARC employs a TCN architecture to process 24 h of continuous physiological input signals. The input tensor is defined as:

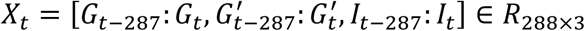

where Gₜ denotes the CGM signal, G′ₜ denotes its temporal derivative, and Iₜ denotes the insulin infusion history. Through a stack of three 1D convolutional layers:

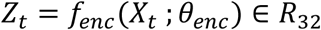

A 32-dimensional physiological latent vector zₜ is generated via a fully connected layer with LayerNorm and SimNorm normalization.

This latent representation is a structured compression that captures meaningful physiological features rather than a simple dimensionality reduction. The policy network combines the current state vector and latent vector zₜ to produce a personalized insulin distribution:

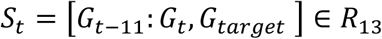

where G_target_ denotes the desired glucose level. An action Aₜ is sampled from the resulting Gaussian distribution using the reparameterization trick:

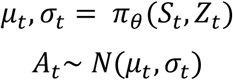

and clipped with a tanh activation to a physiologically valid insulin range of [0.0, 2.0] U. The critic module consists of five Q-networks:

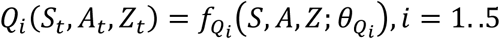

which predict two-hot encoded value distributions. Training stability is ensured via gradient clipping (norm = 10) and soft target updates (τ=0.05).

A key innovation of DPARC is its *cold-start capability*, enabling immediate deployment without any preset user configuration. When fewer than 24 h of data are available, the unobserved portion of the input tensor is initialized after normalization, so padded values represent a neutral normalized input rather than physiologically impossible raw glucose values. With this initialization, the system can enter closed-loop mode after 1 h of observed CGM and insulin records. This approach reduces the initialization burden relative to systems requiring extended run-in or parameter optimization, while allowing early closed-loop operation under limited observed history. It suggests that the latent representation can extract useful physiological context from partially observed states, supporting preclinical feasibility of rapid startup.

DPARC employs Soft Actor-Critic (SAC) (*47*) as its reinforcement learning backbone, selected for its sample efficiency and stable convergence in continuous control tasks. The entropy regularization of SAC naturally promotes exploration during training while maintaining policy stability, which is essential for safe glucose control. The actor network outputs a stochastic policy that balances exploitation of learned strategies with exploration of new physiological conditions, while the ensemble of five Q-networks provides robust value estimation under distributional shifts inherent in cross-individual deployment.

### Modeling physiological variability of in silico environment: Emulating real-world complexity

To train the DPARC under realistic physiological variability rather than a static simulation, we modified the simglucose simulator based on the UVA/Padova model to incorporate intra-day variability scenarios (*48*). Key insulin sensitivity parameters (kp1, kp3, kir, Vmx) were randomly perturbed every 30 min within ±2% of nominal values to simulate real-world factors such as circadian rhythms, stress, and hormonal fluctuations. Meal-specific absorption parameters (f, kabs) were additionally shifted within ±10% at each meal to emulate inter-meal differences in digestion and gastric states (*49*). This approach surpasses simple noise addition by enforcing physiologically meaningful perturbations that rigorously test the adaptive capability of the control algorithm.

For the primary in silico generalization analysis shown in Fig. 2, the meal protocol reflected unpredictable eating behaviors. Three daily meals (07:00, 12:30, 18:00) were jittered uniformly within ±90 min. Carbohydrate amounts were sampled from normal distributions centered at 60, 100, and 80 g for breakfast, lunch, and dinner, respectively. No meal announcements were provided, requiring the controller to detect and respond to postprandial glucose rises solely via CGM. A 10% probability of meal skipping was also introduced so that the evaluation did not rely on a deterministic three-meal schedule. This design departs from traditional meal-announcement paradigms and directly evaluates the system’s capacity to handle unannounced meals in clinical settings.

To maximize policy generalization, 21 virtual training patients were generated via parameter space randomization from 11 original UVA/Padova adults (*35*). Parameters related to insulin demand (Ib, Ith, BW, p2u, kp1–3) were scaled within [-0.3, 0.3], and physiologically plausible samples were selected using Differential Evolution optimization with constraint-based filtering. This ensured the resulting patient cohort exhibited broad yet physiologically consistent variability in insulin sensitivity and carbohydrate absorption profiles.

For controlled mechanistic analyses of latent representation, cold-start convergence, and adaptive response, we employed standardized meal protocols with fixed timing (07:00, 12:30, 18:00) and consistent carbohydrate amounts (40 g, 80 g, 60 g). These fixed-meal experiments were used to isolate latent-state dynamics and physiological perturbation responses from behavioral meal variability, whereas the primary generalization analysis in Fig. 2 used stochastic unannounced meals. This controlled approach enabled systematic evaluation of latent-parameter correlations, cold-start stabilization dynamics, and adaptive responses to physiological perturbations by removing confounding meal variability.

For performance evaluation, we conducted experiments on an extended cohort of 30 virtual patients to assess generalization across diverse physiological profiles. This evaluation cohort comprised 10 patients selected from the original 11 UVA/Padova adults that served as the foundation for training cohort generation, combined with 20 completely independent patients with novel physiological parameters not derived from the original UVA/Padova cohort. The 10 original patients enabled assessment of performance on the base physiological profiles underlying the training data augmentation, while the 20 independent patients provided rigorous evaluation of zero-shot generalization capability to entirely unseen physiological characteristics and parameter ranges. This evaluation strategy ensures comprehensive assessment across both familiar and novel parameter spaces while maintaining statistical power for performance comparisons.

### Pharmacodynamics-informed reward shaping: Learning delayed effects

The delayed effect of insulin is a major challenge in glucose control. While prior RL approaches have addressed this using MDP formulations or auxiliary prediction modules, we introduce a method that directly embeds pharmacodynamic knowledge into the reward structure.

Per-step rewards are defined as follows: a base reward up to 0.5 is granted within ±10 mg/dL of the target glucose range. Penalties of −0.4 to −1.0 are applied for rapid hypoglycemic declines (CGM derivative < −10, −15, −20 mg/dL). The final reward is computed using pharmacodynamic-based temporal weighting over the past 5 h (60 steps):

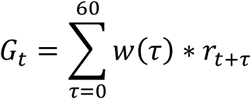

Here, w(τ) represents the insulin action curve derived from the Ordinary Differential Equation-based simulator under basal and bolus conditions. This reward structure provides physiologically meaningful credit assignment, guiding the policy to naturally learn insulin’s delayed metabolic effects.

### Large-scale parallel training: Balancing diversity and efficiency

DPARC was trained over 400+ episodes for 21 virtual patients. Using the Ray framework, a separate actor was executed for each patient in parallel. This design improved computational efficiency while preserving each patient’s physiological distinctiveness during training.

In conventional mixed-training schemes, experience from multiple patients is aggregated into a single replay buffer, risking dilution of patient-specific dynamics. Our approach maintained each patient’s trajectory in an independent buffer and employed a 60-step sliding window to collect trajectories over 24-h episodes. Mini-batches (size 256) were sampled independently from each buffer, allowing the algorithm to retain personalized learning signals across patients. This architecture ensures the policy can generalize across diverse patient groups without overfitting to an average patient, a critical requirement for real-world deployment.

The model was updated over 576 steps per episode, experimentally chosen to ensure convergence for complex physiological dynamics. Every 10 episodes, performance was evaluated over 5 days on both 10 training and 20 unseen test patients. This frequent validation enabled early detection of overfitting and selection of optimal checkpoints for deployment.

### In vivo validation: Bridging simulation and biology

To evaluate the robustness of DPARC against rapid physiological dynamics, we first performed closed-loop control experiments on six male Sprague–Dawley rats (300–500 g) with streptozotocin-induced diabetes (75 mg/kg intraperitoneal injection). All procedures were approved by the Institutional Animal Care and Use Committee of Daegu Haany University (IACUC number: DHU2025-030). The autonomous system comprised a Dexcom G7 CGM and a Dana-i insulin pump (delivering 1:10 diluted Humalog) controlled via the CloudLoop smartphone application. DPARC was deployed directly (zero-shot) without any pre-tuning or open-loop warm-up. Rats were fed standardized chow (5–10 g) three times daily without meal announcements, and data from all six subjects were included in the analysis.

Subsequently, to assess clinical translational potential, fully closed-loop experiments were conducted on five Yorkshire swine (20–40 kg) with streptozotocin-induced diabetes (150 mg/kg via ear vein injection). All procedures were approved by the K-MEDI hub IACUC (approval number: KMEDI-23051103-00). The setup utilized the same hardware ecosystem (Dexcom G7, Dana-i with Novorapid and Humalog). For DPARC evaluation (n=5), the controller initiated closed-loop control after only 1 hour of open-loop monitoring without parameter configuration. Meals (200–300 g) were delivered via automated feeders without announcement. To benchmark performance, a crossover study was conducted in a subset of animals (n=2) comparing DPARC against the AAPS. The crossover protocol proceeded as follows: (i) an open-loop run-in period to optimize individual therapy parameters (basal rate, ISF, CR) required for AAPS (specific profiles provided in Supplementary Table S6); (ii) the AAPS closed-loop session; (iii) an open-loop washout period; and (iv) the DPARC session. Diabetic status (hyperglycemia >200 mg/dL) was confirmed prior to each experimental session and verified again upon study completion. While AAPS operated using the optimized profile derived from the run-in period, DPARC operated in a strictly zero-shot manner without prior information.

## List of Supplementary Materials

Supplementary Notes 1-4

Supplementary Figures. S1-S11

Supplementary Tables S1-S6

## Acknowledgments

This research was supported by the University Technology Commercialization Promotion Program through the Commercializations Promotion Agency for R&D Outcomes (COMPA) funded by the National Research Foundation of Korea (NRF) (RS-2024-00426901); the National Research Foundation of Korea (NRF) grant funded by the Korea government (MSIT) (RS-2025-00517742); the Korea Health Industry Development Institute (KHIDI) (HI23C004800); and KHIDI (RS-2023-00215716).

## Author contributions

Junyoung Yoo: Conceptualization, Methodology, Software, Formal analysis, Investigation, Data curation, Writing original draft, Writing review & editing, Visualization. Vega Pradana Rachim: Methodology, Software. Yein Lee: Software. Jaeyeon Lee: Methodology, Supervision, Project administration. Sung-Min Park: Methodology, Project administration, Supervision, Writing review & editing.

## Data and materials availability

All data needed to evaluate the conclusions in the paper are present in the paper and/or the Supplementary Materials.

