## Supplementary Information for "Zero-shot automated insulin delivery for type 1 diabetes via dynamic physiology-aware reinforcement learning"

Junyoung Yoo *et al.*

*Corresponding author.
Jaeyeon Lee:
Sung-Min Park:

**This PDF file includes:**

Supplementary Notes 1-4
Figs. S1-S11
Tables S1-S6
Reference (35)

**Supplementary Text**

**Supplementary Note 1.** DPARC pseudo code

**Input:**

Patient sets (train, test),
Environment $E$ (Type 1 diabetes (T1D simulator),
Replay buffers $D$,
Agent $\pi_\theta$ (DPARC with parameters $\theta$)

**Output:**

Trained policy $\pi_\theta$ for BG control

```
Initialize DPARC agent π_θ and replay buffers D
for episode = 1 to N do
    if episode = 1 then
        set exploration ← random actions
    else
        set exploration ← policy actions
    end if

    for each patient p in train set do
        Launch parallel Actor(p)
        τ_p ← CollectTrajectory(E, π_θ, exploration)
        Store τ_p into corresponding replay buffer D_p
    end for

    for update step = 1 to U do
        Sample minibatch B from each D_p
        Update critic Q_φ using Bellman backup
        Update actor π_θ using policy gradient
        Update temperature α for entropy regularization
    end for

    if episode mod 5 = 0 then
        Save current model parameters θ
    end if

    if episode mod 10 = 0 then
        for each patient q in test set do
```

```
            Evaluate policy πθ on E(q)
            Record metrics: TIR, TAR, TBR, survival
            Save CGM/insulin trajectories
        end for
    end if
end for
```

**Supplementary Note 2.** Rodent data acquisition

Six Sprague–Dawley rats (300–500 g) were employed to evaluate the cold-start performance of DPARC. All animal procedures were approved by the Institutional Animal Care and Use Committee (IACUC number: DHU2025-030) and conducted in accordance with established guidelines for the care and use of laboratory animals.

Diabetes was induced by a single intraperitoneal injection of streptozotocin (75 mg/kg; Sigma–Aldrich) as described in previous studies, and onset was confirmed when fasting blood glucose exceeded 200 mg/dL. For closed-loop studies, the same experimental setup as in swine was applied: Dexcom G7 continuous glucose monitoring sensors, Dana-i insulin pumps delivering Humalog insulin (1:10 diluted with saline), and the CloudLoop application implementing the DPARC policy via xDrip+ data relay.

Rats were provided standardized chow in controlled portions of 5–10 g per meal, administered three times daily. Each animal underwent fully closed-loop testing under DPARC without any preconditioning or subject selection. Cold-start evaluation was performed directly at deployment, and all recorded data were included in the analysis without exclusion.

**Supplementary Note 3.** Swine data acquisition

Five Yorkshire swine (20–40 kg) were employed for closed-loop validation of DPARC. All procedures were approved by the K-MEDI hub Institutional Animal Care and Use Committee (IACUC number: KMEDI-23051103-00) and conducted in accordance with established guidelines for the care and use of laboratory animals. Additionally, routine lighting cycle and standard room temperatures were maintained. Pigs were housed in an air-conditioned room with a 12 hr light-dark cycle and controlled temperature (24 ± 2°C). The animals were premedicated with atropine sulfate (Atropine Sulfate®, Huons Co., Ltd., 0.04 mg/kg, IM) and xylazine hydrochloride (Rompun®, Bayer Co., Ltd., 4.4 mg/kg, IM) for immobilization and tracheal intubation. The anesthesia was maintained with 2% isoflurane under pure oxygen. Diabetes was induced by intravenous administration of streptozotocin (150 mg/kg; Sigma–Aldrich S0130), and onset was confirmed when fasting blood glucose exceeded 200 mg/dL. All animals were allowed to recover from anesthesia following the completion of streptozotocin administration and were continuously monitored with supplemental oxygen support until stable anesthetic recovery was achieved. Heart rate, respiratory rate, and blood pressure were continuously monitored using physiological monitoring equipment throughout the entire anesthesia and drug administration procedures.

For the closed-loop study, swine were equipped with Dexcom G7 continuous glucose monitoring sensors and Dana-i insulin pumps delivering Novorapid and Humalog insulin. CGM data were streamed to a smartphone via the xDrip+ open-source application, and insulin dosing was determined in real time by the CloudLoop application implementing the DPARC policy. CloudLoop*(35)* directly communicated dosing commands to the Dana-i pump, serving as the central hub for data integration and control. The deployed policy was identical to the in silico trained model, with no retraining or subject-specific parameterization.

For the subset of animals (Subjects 4 and 5) engaged in the crossover benchmark against Android Artificial Pancreas System (AAPS), specific clinical parameters were determined during an open-loop run-in period to configure the AAPS controller; see Supplementary Table S6 for detailed profiles. Note that these parameters were used exclusively for AAPS and were not provided to DPARC.

Due to species-specific behaviors, uninterrupted multi-day data collection in swine was not feasible. In several trials, sensor detachment, cannula line damage, feeder malfunction, or pump destruction (chewing by the swine) led to incomplete records, making those segments unsuitable for analysis. For stability and safety, both the infusion cannula and CGM sensor were placed on the dorsal side of the swine; however, the thickness and mechanical properties of porcine skin and subcutaneous tissue in this region made consistent subcutaneous insertion challenging. This limitation likely introduced variability in both sensor accuracy and insulin delivery reliability. Consequently, we report results from the longest consecutive segments where continuous sensor and infusion data were available. During the experimental period, swine were fed standardized chow meals three times daily. Postprandial monitoring windows were defined as 5 h after each meal, and overnight monitoring was conducted from 1:00–7:00 a.m.

**Supplementary Note 4.** Safety constraints and fail-safe mechanisms

To ensure safe operation during preclinical validation, multiple layers of safety constraints were implemented within the CloudLoop application and DPARC algorithm architecture.

**Hardware-level safety measures:**

- Minimum time interval between bolus deliveries was enforced at 5 minutes
- CGM sensor failure detection triggered automatic suspension of insulin delivery

- Pump error resulted in immediate cessation of control algorithm

**Algorithm-level safety constraints:**

- Insulin action clipping with tanh activation limited individual bolus doses to 0.0–2.0 U range.
- Communication timeout between CGM and pump (>30 min) resulted in algorithm suspension

**Monitoring and intervention protocols:**

- Continuous veterinary supervision was maintained throughout all experimental sessions
- Emergency glucose administration was immediately available for prolonged or clinically concerning hypoglycemic episodes.

These multi-layered safety measures ensured that preclinical validation maintained animal welfare as the highest priority while enabling robust evaluation of DPARC performance under controlled conditions.

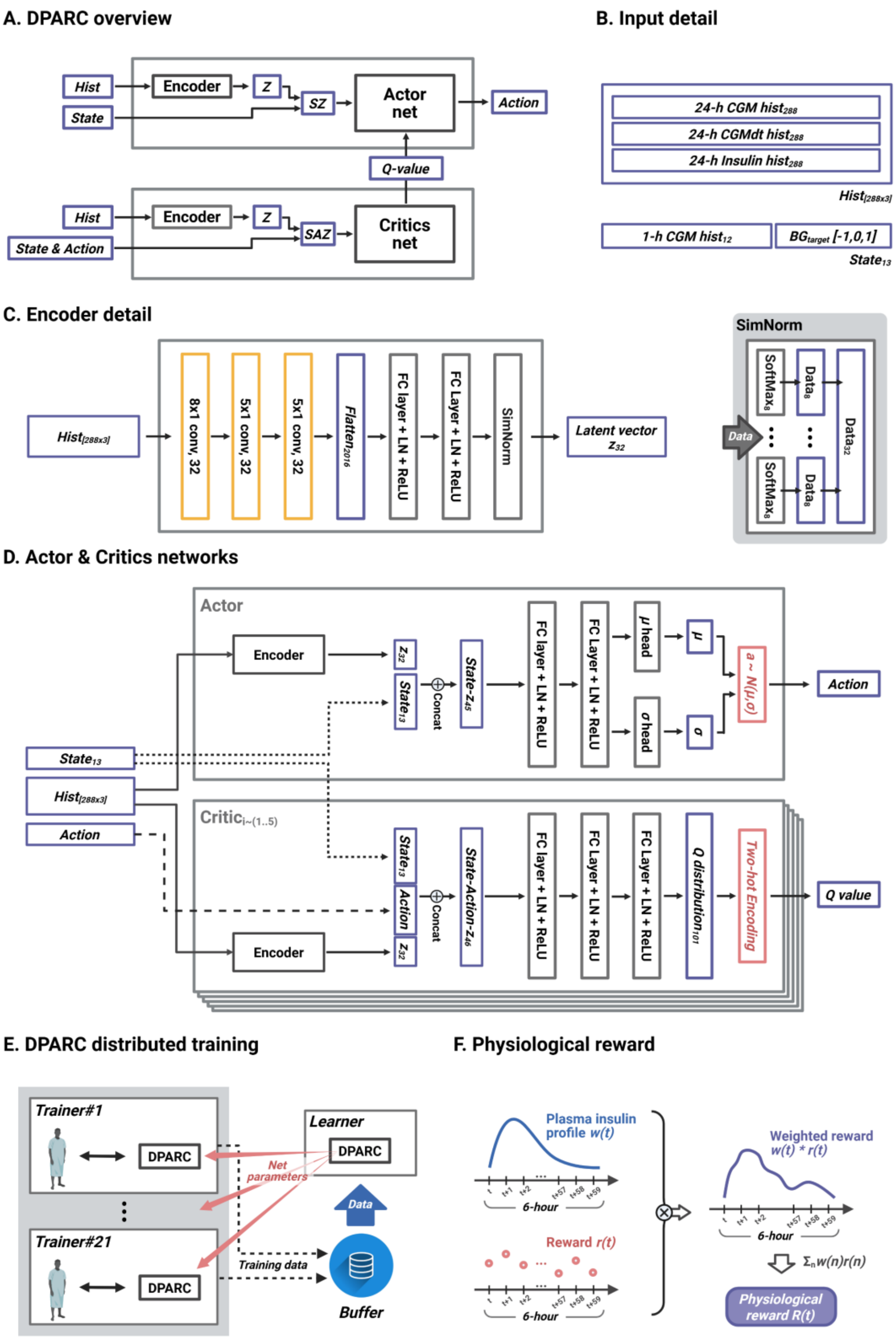


**Fig. S1. DPARC architecture overview and training framework. (A)** System overview showing the dual-pathway architecture with shared temporal encoder. The encoder processes 24-

h history to generate latent vector z, which conditions both the actor network (for insulin delivery actions) and ensemble of critic networks (for Q-value estimation). **(B)** Input specifications detailing the temporal structure of physiological data. The encoder receives 24-h histories of CGM readings, CGM rate of change (CGMdt), and insulin delivery records ($Hist_{288\times3}$), while the actor network processes current state information including recent 1-h CGM history and target glucose level ($State_{13}$). **(C)** Temporal convolutional encoder architecture comprising three Conv1D layers with increasing receptive fields, followed by fully connected layers with LayerNorm and SimNorm for latent representation learning. The encoder compresses 288 timesteps of multivariate physiological data into a 32-dimensional latent vector z. **(D)** Actor and critic network architectures. The actor network combines state and latent representations to output continuous insulin actions via parameterized Gaussian distributions with tanh activation bounds. The critic ensemble consists of five Q-networks processing state-action-latent concatenations to estimate value distributions using two-hot encoding. **(E)** Distributed training framework employing parallel trainers across 21 virtual patients, with independent replay buffers maintaining patient-specific trajectories while enabling generalized policy learning. **(F)** Physiological reward shaping incorporating insulin pharmacodynamics through temporal weighting. The reward function applies plasma insulin profile w(t) over a 6-h horizon to provide biologically informed credit assignment, aligning policy optimization with insulin action delays.

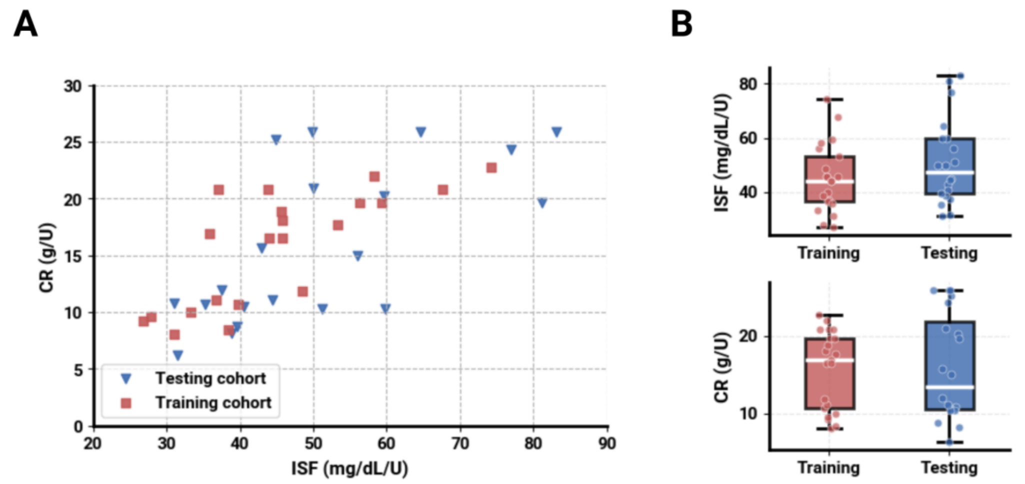


**Fig. S2. Physiological parameter distributions across training and evaluation cohorts. (A)** Scatter plot showing the distribution of insulin sensitivity factor (ISF) and carbohydrate ratio (CR) parameters for training cohort (n=21, red squares) and testing cohort (n=30, blue triangles). The training cohort was generated via parameter space randomization from 11 original UVA/Padova adults with parameter augmentation. The testing cohort comprised 10 patients from the original UVA/Padova cohort plus 20 completely independent patients with novel physiological parameters. **(B)** Box plot distributions of ISF (top) and CR (bottom) parameters with individual data points overlaid. Box plots display median, quartiles, and whiskers extending to 1.5× interquartile range. Individual scatter points show the complete distribution of parameter values across both cohorts.

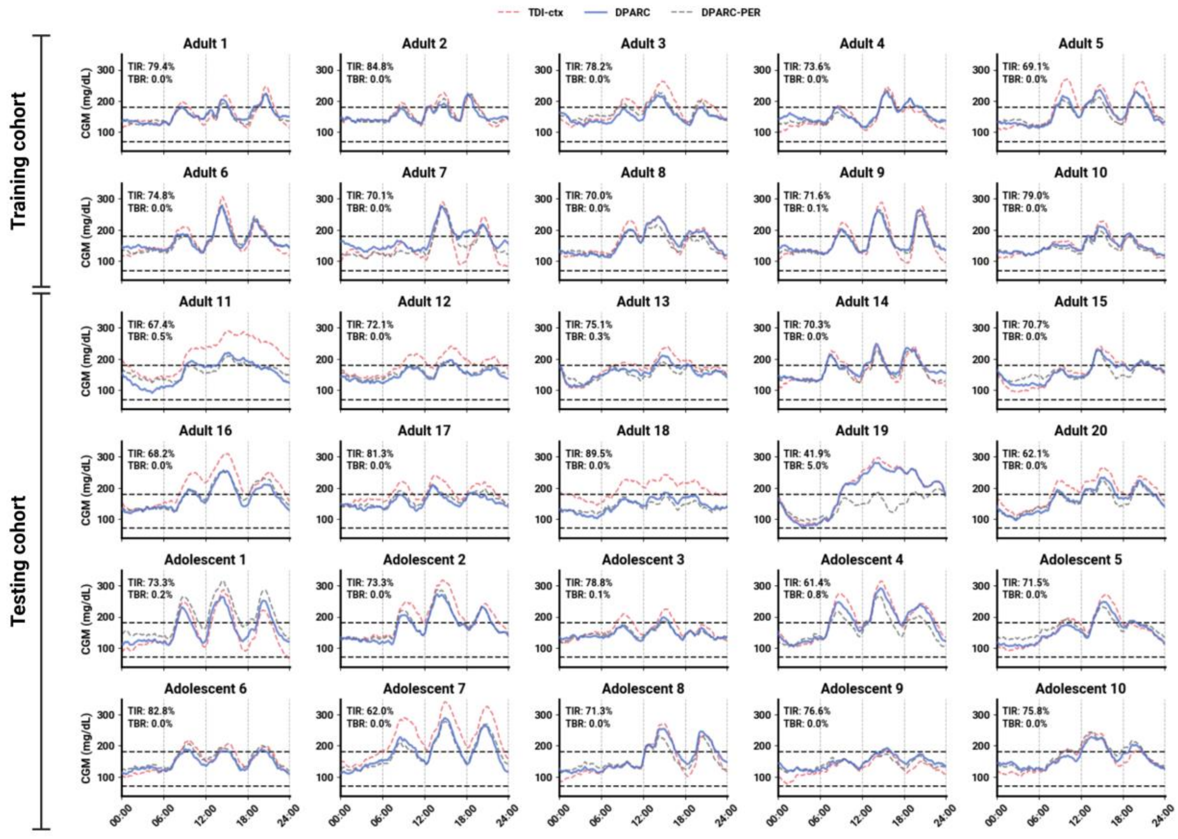


**Fig. S3. In silico mean daily glucose profiles for individual virtual subjects.** Mean continuous glucose monitoring (CGM) profiles over 24-h for each virtual subject. For every individual, 10 consecutive simulation days were stacked and averaged to yield a representative daily CGM trace. Results are shown for the training cohort (Adults 1–10) and testing cohort (Adults 11–20, Adolescents 1–10) under three controllers: TDI-context (TDI-ctx) (red dashed), the proposed DPARC (blue solid), and DPARC-personalized (DPARC-PER) (gray dashed). Reported time in range (TIR) and time below range (TBR) values correspond to the averaged outcomes over the 10-day period.

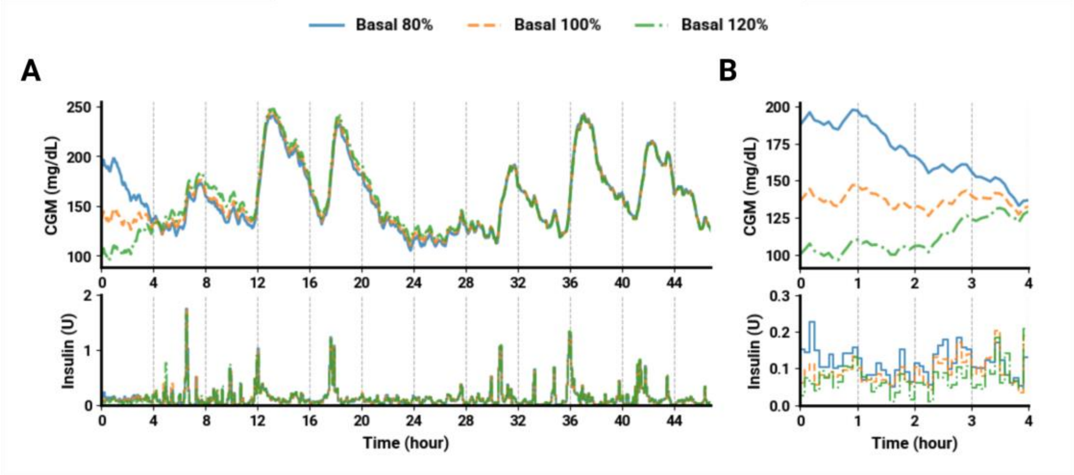


**Fig. S4. Cold-start behavior under perturbed basal infusion.** Mean simulated glucose–insulin dynamics over 48-h in 30 in silico patients from the UVA/Padova cohort. A controlled fixed-meal protocol was used to isolate cold-start behavior from meal-pattern variability and included three daily meals (40 g breakfast, 80 g lunch, 60 g dinner). Basal infusion was set to 80%, 100%, or 120% of the optimal basal rate to induce diverse initial glycemic states. **(A)** Mean CGM and insulin traces over two simulation days, showing DPARC performance after cold start conditions. **(B)** Mean CGM and insulin traces during the first 4 h after initialization, highlighting controller behavior under unstable starting glucose levels.

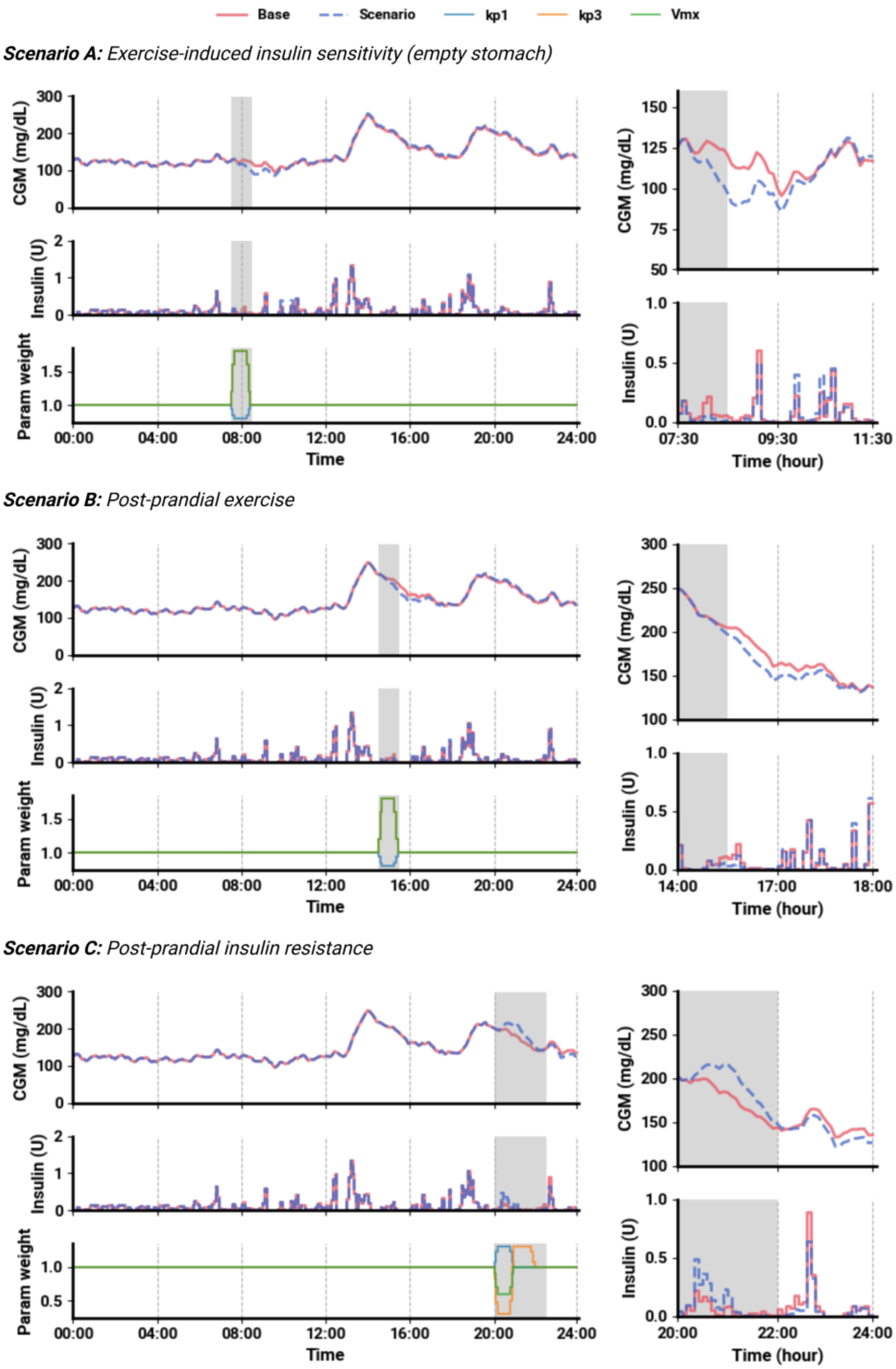


**Fig. S5. Simulated exercise and insulin resistance scenarios modeled by transient perturbations of physiological parameters.** Closed-loop simulations were performed under three conditions with meals at lunch (80 g CHO) and dinner (60 g CHO). **(A)** Exercise-induced insulin sensitivity in the fasting state (07:30, no meal). Transient increases in insulin sensitivity (kp3), peripheral utilization (Vmx), and reduced endogenous glucose production (kp1) led to lower post-exercise glucose and reduced insulin dosing. **(B)** Post-prandial exercise (14:30, following 80 g CHO). Similar parameter modulations after lunch reduced the post-prandial glucose excursion. **(C)** Post-prandial insulin resistance (20:00, following 60 g CHO). Temporary decreases in kp3 and Vmx with increased kp1 produced elevated post-prandial glucose levels and higher insulin demand. Parameter weight traces (bottom panels) indicate the timing and magnitude of perturbations (grey shading). Right panels show magnified views of exercise/insulin resistance periods.

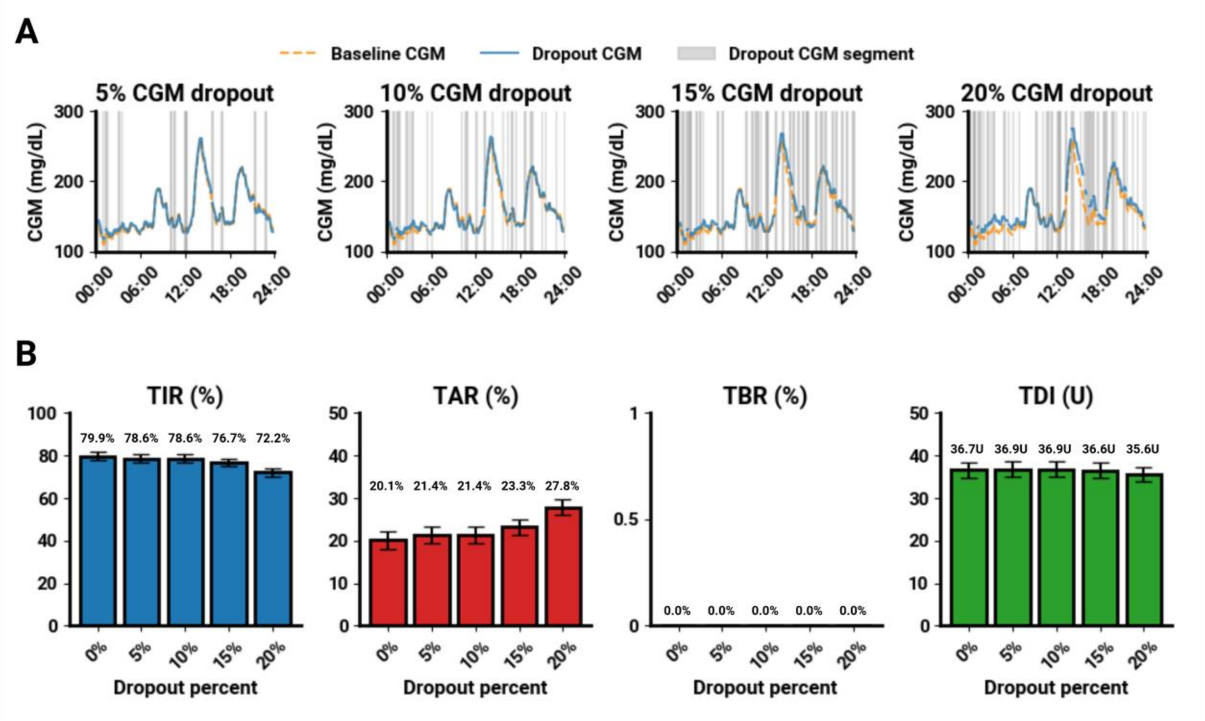


**Fig. S6. Sensitivity of DPARC to random CGM dropout.** 3-day simulations were conducted in 30 in silico adults from the UVA/Padova cohort under a fixed meal protocol (40 g breakfast, 80 g lunch, 60 g dinner). During simulation, CGM signals were randomly removed at dropout rates of 0%, 5%, 10%, 15%, and 20%. **(A)** Mean CGM trajectories with dropout segments highlighted for different dropout rates. **(B)** Summary performance metrics including TIR, time above range (TAR), TBR, and TDI as a function of dropout rate. DPARC preserved overall TIR with low hypoglycemia exposure under the tested dropout conditions. Results are shown as mean ± SEM.

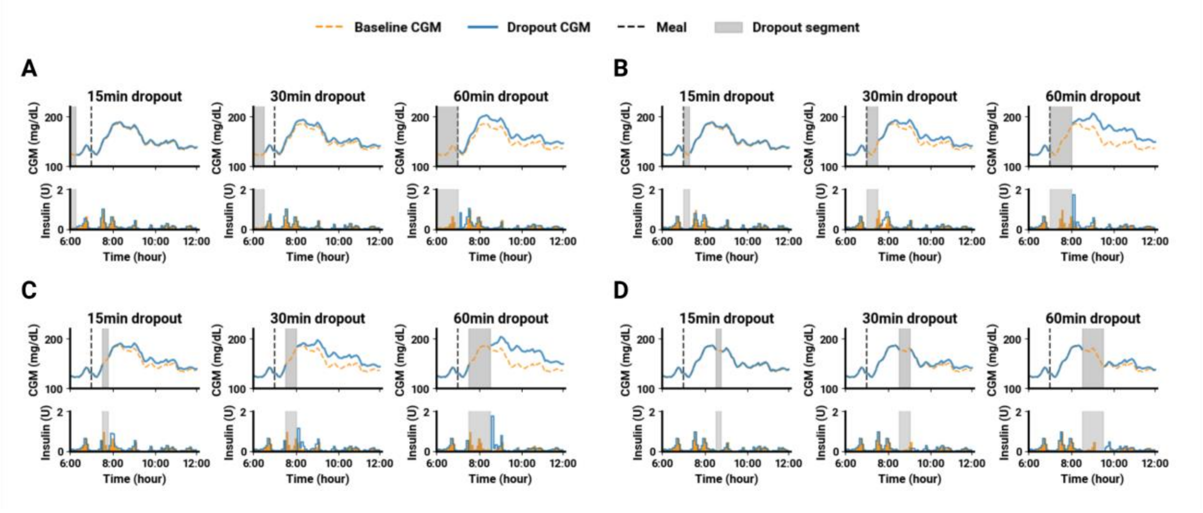


**Fig. S7. Robustness to meal-phase CGM dropout.** Six-hour simulations in 30 in silico adults from the UVA/Padova cohort under the fixed meal protocol (40 g breakfast, 80 g lunch, 60 g dinner). CGM data were removed for 15, 30, or 60 min during four distinct phases of the meal response: **(A)** 1-h before meal onset, **(B)** at meal initiation, **(C)** 30-min postprandially during the glucose rise, and **(D)** 90-min postprandially during the glucose decline. DPARC maintained closed-loop operation under the tested dropout scenarios. Results represent mean CGM trace.

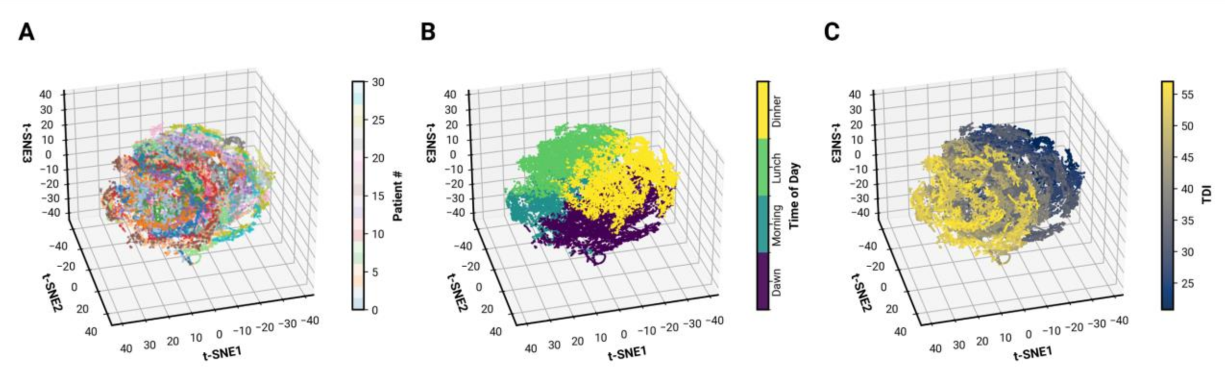


**Fig. S8. Latent space embeddings of DPARC encoder.** Three-dimensional t-SNE projections of the 32-dimensional latent vector z, inferred every 5 min from 24-h CGM and insulin histories in 30 virtual patients under fixed meal conditions (40 g breakfast, 80 g lunch, 60 g dinner; no intraday IS drift). **(A)** Color coding by patient ID shows partial overlap across individuals, indicating that the latent space does not encode subject identity as a dominant factor. **(B)** Coloring by time of day illustrates temporal organization of the latent space under the controlled fixed-meal protocol; this analysis was used for mechanistic visualization rather than for meal-pattern generalization testing. **(C)** A smooth gradient appears when colored by total daily insulin (TDI), suggesting that insulin demand is embedded as a continuous physiological axis. Together, these visualizations illustrate that the latent representation captures physiologically relevant structure (daily cycles, insulin demand) while remaining broadly generalizable across patients.

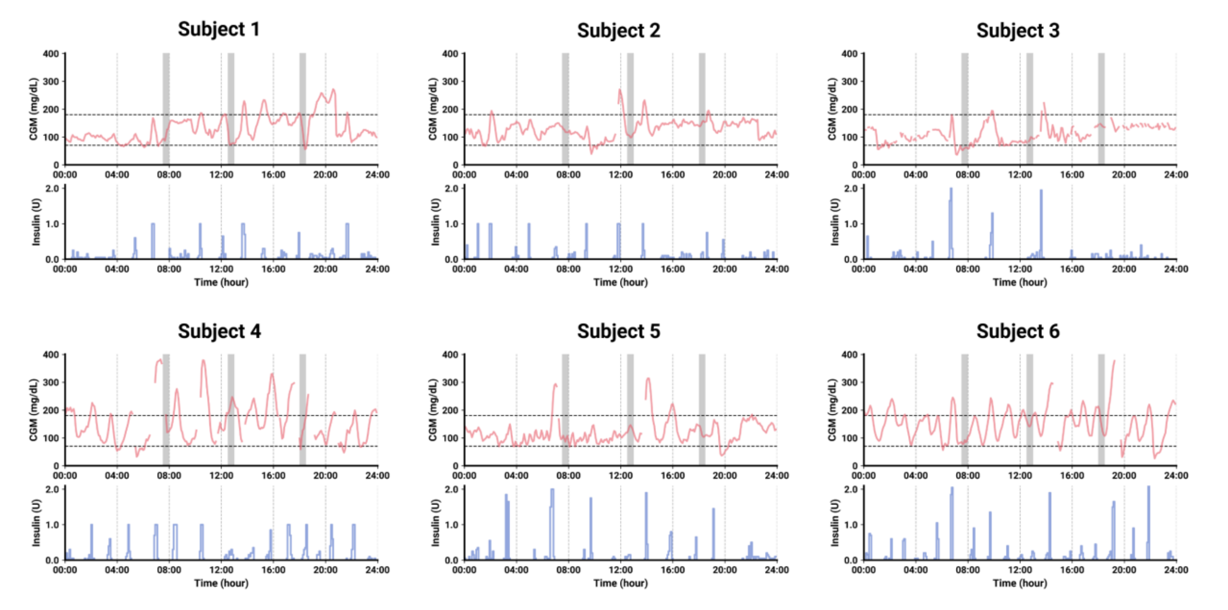


**Fig. S9. Individual Rat closed-loop traces under DPARC control.** Continuous glucose monitoring (CGM, red) and insulin delivery (blue) are shown for six diabetic rats (Rat 1–6). Dashed horizontal lines denote glycemic thresholds of 70 and 180 mg/dL. Despite inter-animal variability in glucose excursions and insulin requirements, DPARC executed zero-shot closed-loop control across sessions without algorithmic modification or subject-specific tuning.

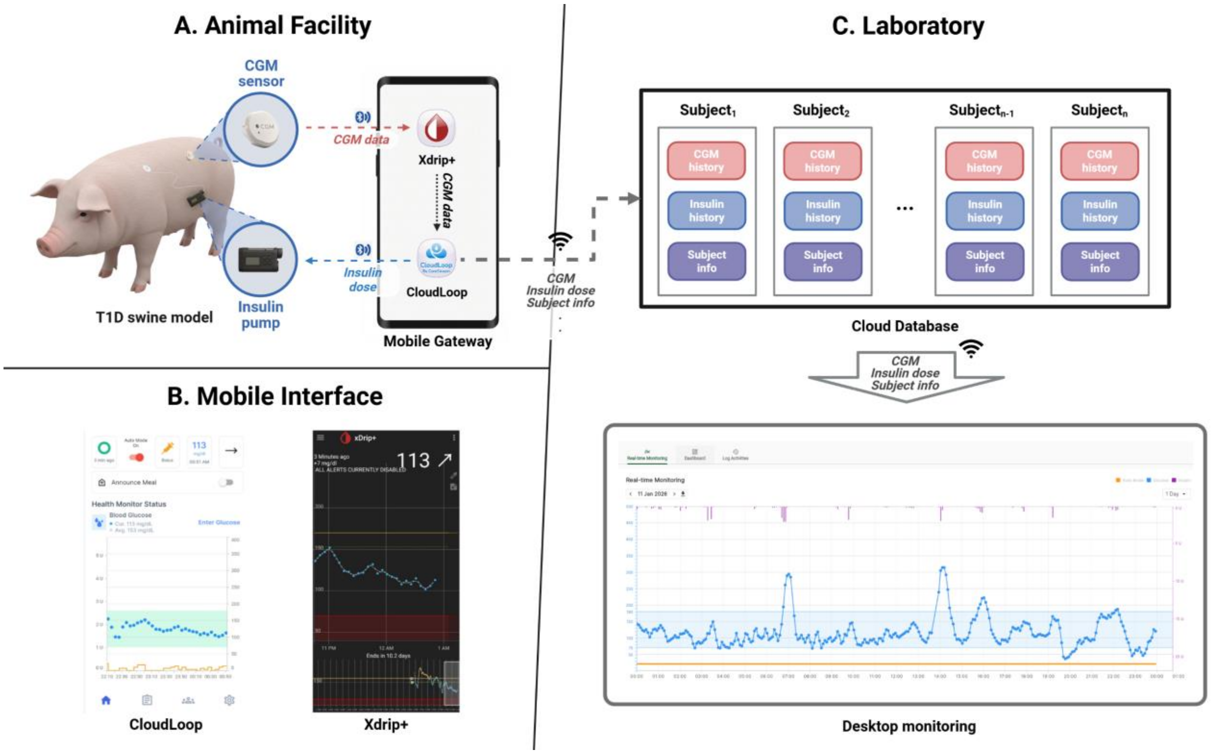


**Fig. S10. Schematic illustration of the pre-clinical data acquisition and monitoring platform. (A)** The experimental setup for the T1D swine model. The subject is equipped with a Dexcom G7 CGM sensor and a Dana-i insulin pump, which communicate via Bluetooth with a smartphone gateway. **(B)** The smartphone runs the Xdrip+ application for CGM data collection and the CloudLoop application for implementing the DPARC policy and issuing insulin commands. **(C)** The remote monitoring infrastructure (Laboratory). Real-time physiological data (CGM, insulin history) are synchronized from the smartphone to a central database, enabling researchers to monitor the glycemic status of multiple subjects simultaneously via a desktop dashboard.

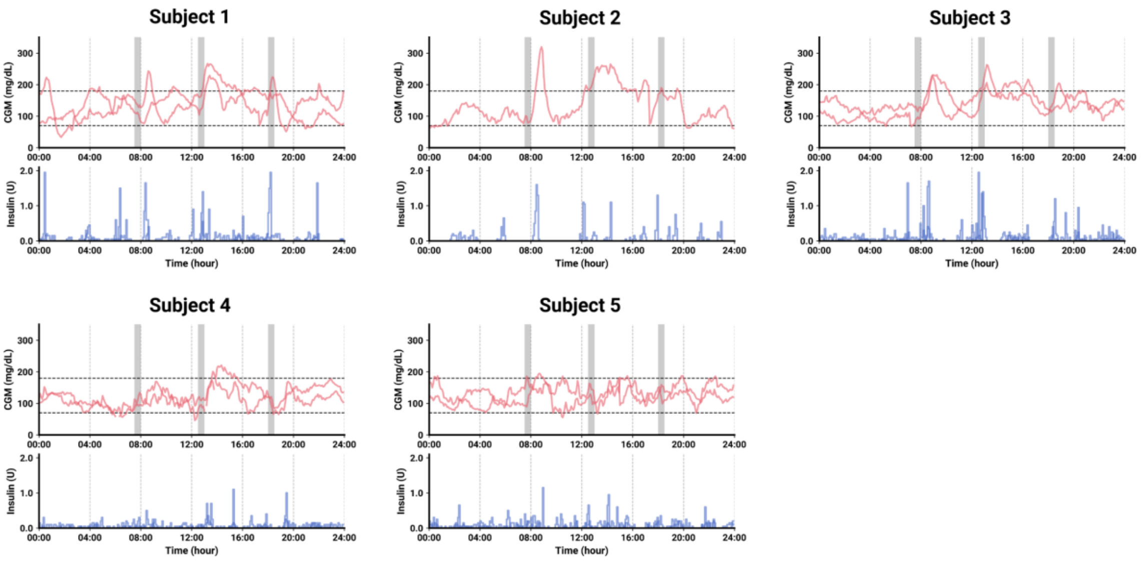


**Fig. S11. Individual swine closed-loop traces under DPARC control.** Continuous glucose monitoring (CGM, red) and insulin delivery (blue) are shown for five diabetic Yorkshire swine (Swine 1–5) across two experimental sessions. Grey shaded areas indicate meal times (breakfast, lunch, and dinner). Dashed horizontal lines denote glycemic thresholds of 70 and 180 mg/dL. Despite inter-animal variability in glucose excursions and insulin requirements, DPARC maintained stable glucose control across sessions without algorithmic modification or subject-specific tuning.

**Table S1. DPARC neural network architecture specifications.** Detailed configuration of the Dynamic Physiology-Aware RL Controller components. The temporal encoder employs a three-layer convolutional architecture followed by fully connected layers to compress 24-h physiological histories into latent representations. The actor network combines current state and latent features to generate continuous insulin delivery actions through parameterized Gaussian distributions. The critic ensemble consists of five Q-networks that process state-action-latent concatenations for robust value estimation using distributional reinforcement learning with two-hot encoding. Training employs Soft Actor-Critic (SAC) with entropy regularization, gradient clipping, and soft target updates for stable learning. Safety constraints include action bounds, log-standard deviation limits, and gradient norm clipping to ensure safe insulin delivery during deployment.

| Component | Layer/Module | Configuration |
|---|---|---|
| **Temporal Encoder** | Conv1D Layer 1 | in_channels=3, out_channels=32, kernel_size=8, stride=4 |
| | Activation | LeakyReLU |
| | Conv1D Layer 2 | in_channels=32, out_channels=32, kernel_size=5, stride=1 |
| | Activation | LeakyReLU |
| | Conv1D Layer 3 | in_channels=32, out_channels=32, kernel_size=5, stride=1 |
| | Activation | LeakyReLU |
| | Flatten | - |
| | Linear Layer 1 | in_features=2016, out_features=256 |
| | Normalization | LayerNorm(256) |
| | Activation | ReLU |
| | Linear Layer 2 | in_features=256, out_features=32 |
| | Normalization | LayerNorm(32) + SimNorm |
| **Actor Network** | Shared MLP | input_dim=state_dim+32, hidden=[256,256], activation=ReLU |
| | Mean Layer | Linear(256 → action_dim) |
| | Std Layer | Linear(256 → action_dim) |
| | Action Bounds | tanh activation, max_action scaling |
| **Q-Network Ensemble** | Shared Encoder | Same as Temporal Encoder |
| | Q-Networks (×5) | MLP(state_dim+action_dim+32 → 101, hidden=[256,256,256]) |
| | Output | Two-hot encoding for value distribution (bins=101) |
| **Training Configuration** | Optimizer | Adam (lr=3e-4) |
| | Entropy Target | -1 |
| | Discount Factor | γ=0.98 |
| | Soft Update | τ=0.05 |
| | Gradient Clipping | max_norm=10 |
| | Batch Size | 256 |

| | | |
|---|---|---|
| **Input Specifications** | History Window | 288 timesteps (24h × 12 samples/h) |
| | Input Channels | 3 (CGM, CGMdt, Insulin) |
| | State Vector | Current CGM + target glucose |
| | Action Space | Continuous insulin delivery [0.0, 2.0] U |
| **Safety Constraints** | Action Clipping | tanh(μ) |
| | Log-std Bounds | Clamped to [-20, 2] |
| | Gradient Clipping | L2 norm ≤ 10 |

**Table S2. In silico performance (mean ± SD) across 30 virtual patients.** Values represent the mean outcome of five independent trials per subject. Statistical testing was performed with repeated-measures ANOVA across algorithms (TDI-ctx, DPARC, DPARC-PER), followed by FDR-adjusted paired comparisons. Stars indicate significant difference versus TDI*: $p < 0.05$* (*)*,* $p < 0.01$ (**)*,* $p < 0.001$ (***); ns = not significant. DPARC-PER corresponds to a personalized per-subject model and is reported as an upper-bound benchmark. DPARC, trained in a context-free manner without patient-specific inputs, significantly outperformed the TDI-ctx comparator and achieved performance approaching the PER upper bound across key metrics.

| | **TDI-ctx** | **DPARC** | **DPARC-PER** |
|---|---|---|---|
| **$TIR_{70-180}$ (%)** | 63.4 ± 12.1 | 72.5 ± 8.5** | 74.8 ± 6.9*** |
| **$TIR_{70-140}$ (%)** | 42.1 ± 15.1 | 43.7 ± 7.9 (ns) | 45.8 ± 8.3 (ns) |
| **$TAR_{180}$ (%)** | 35.2 ± 12.3 | 27.2 ± 7.9** | 24.7 ± 6.2*** |
| **$TAR_{250}$ (%)** | 11.6 ± 7.6 | 6.3 ± 5.4*** | 5.3 ± 4.7*** |
| **$TBR_{70}$ (%)** | 1.4 ± 2.6 | 0.2 ± 0.9 (ns) | 0.5 ± 1.9 (ns) |
| **$TBR_{54}$ (%)** | 0.6 ± 1.4 | 0.1 ± 0.3 (ns) | 0.3 ± 1.6 (ns) |
| **Mean BG (mg/dL)** | 168 ± 19 | 160 ± 10 (ns) | 158 ± 8* |
| **SD (mg/dL)** | 57 ± 13 | 47 ± 11*** | 44 ± 11*** |
| **CV (%)** | 34.4 ± 7.4 | 29.0 ± 5.4*** | 28.1 ± 6.2*** |
| **TDI (U)** | 37.3 ± 9.9 | 38.8 ± 11.1 (ns) | 39.3 ± 11.6* |

**Abbreviations: $TIR_{70-180}$**, Time In Range (70–180 mg/dL); **$TIR_{70-140}$**, Time In Range (70–140 mg/dL); **$TAR_{180}$**, Time Above Range (>180 mg/dL); **$TAR_{250}$**, Time Above Range (>250 mg/dL); **$TBR_{70}$**, Time Below Range (<70 mg/dL); **$TBR_{54}$**, Time Below Range (<54 mg/dL); **BG**, Blood Glucose; **SD**, Standard Deviation; **CV**, Coefficient of Variation; **TDI**, Total Daily Insulin.

**Note**: TDI-ctx refers to the baseline control algorithm. DPARC denotes the proposed context-free model, and DPARC-PER denotes the personalized model.

**Table S3. Individual rat closed-loop performance (mean ± SD) under DPARC control.** Results from six streptozotocin-induced diabetic Sprague–Dawley rats tested under 24-h closed-loop control.

| | $TIR_{70\text{-}180}$ (%) | $TIR_{70\text{-}140}$ (%) | $TAR_{180}$ (%) | $TAR_{250}$ (%) | $TBR_{70}$ (%) | $TBR_{54}$ (%) | Mean BG (mg/dL) | SD (mg/dL) | CV (%) |
|---|---|---|---|---|---|---|---|---|---|
| **Subject 1** | 91.3 | 56.3 | 5.2 | 0.7 | 3.5 | 0.7 | 129 | 34 | 26.0 |
| **Subject 2** | 83.0 | 58.3 | 14.2 | 2.1 | 2.8 | 0.0 | 131 | 47 | 36.3 |
| **Subject 3** | 56.7 | 43.3 | 35.2 | 11.1 | 8.0 | 1.9 | 157 | 76 | 48.2 |
| **Subject 4** | 88.4 | 75.7 | 2.0 | 0.0 | 9.6 | 1.6 | 110 | 32 | 29.0 |
| **Subject 5** | 67.4 | 37.3 | 27.6 | 2.9 | 5.0 | 3.6 | 151 | 57 | 37.5 |
| **Subject 6** | 88.7 | 72.2 | 6.7 | 2.8 | 4.6 | 1.8 | 120 | 44 | 36.9 |
| **Total** | **79.3 ± 12.8** | **57.2 ± 13.9** | **15.2 ± 12.3** | **3.3 ± 3.7** | **5.6 ± 2.4** | **1.6 ± 1.1** | **133 ± 16** | **48 ± 15** | **35.7 ± 7.1** |

**Abbreviations:** **$TIR_{70\text{-}180}$**, Time In Range (70–180 mg/dL); **$TIR_{70\text{-}140}$**, Time In Range (70–140 mg/dL); **$TAR_{180}$**, Time Above Range (>180 mg/dL); **$TAR_{250}$**, Time Above Range (>250 mg/dL); **$TBR_{70}$**, Time Below Range (<70 mg/dL); **$TBR_{54}$**, Time Below Range (<54 mg/dL); **BG**, Blood Glucose; **SD**, Standard Deviation; **CV**, Coefficient of Variation

**Table S4. Individual swine closed-loop performance (mean ± SD) under DPARC control**. Performance metrics are reported for nine sessions across five diabetic Yorkshire swine. All trials were conducted under an unannounced three-meal protocol without algorithm modification or subject-specific tuning.

| | $TIR_{70\text{-}180}$ (%) | $TIR_{70\text{-}140}$ (%) | $TAR_{180}$ (%) | $TAR_{250}$ (%) | $TBR_{70}$ (%) | $TBR_{54}$ (%) | Mean BG (mg/dL) | SD (mg/dL) | CV (%) |
|---|---|---|---|---|---|---|---|---|---|
| **Subject 1 (Day 1)** | 86.5 | 59.0 | 11.1 | 0.0 | 2.4 | 0.0 | 128 | 39 | 30.4 |
| **Subject 1 (Day 2)** | 71.2 | 37.2 | 21.5 | 2.4 | 7.3 | 3.1 | 146 | 49 | 33.4 |
| **Subject 2 (Day 1)** | 70.8 | 58.3 | 22.9 | 5.9 | 6.2 | 0.0 | 135 | 57 | 41.8 |
| **Subject 3 (Day 1)** | 89.9 | 62.8 | 10.1 | 0.7 | 0.0 | 0.0 | 137 | 34 | 25.2 |
| **Subject 3 (Day 2)** | 79.9 | 51.7 | 19.4 | 0.0 | 0.7 | 0.0 | 141 | 40 | 28.3 |
| **Subject 4 (Day 1)** | 89.9 | 61.9 | 7.0 | 0.0 | 3.1 | 0.7 | 124 | 37 | 30.1 |
| **Subject 4 (Day 2)** | 94.4 | 83.9 | 1.4 | 0.0 | 4.2 | 0.4 | 114 | 24 | 21.5 |
| **Subject 5 (Day 1)** | 94.4 | 54.5 | 4.9 | 0.0 | 0.7 | 0.0 | 135 | 28 | 20.9 |
| **Subject 5 (Day 2)** | 96.9 | 79.9 | 1.7 | 0.0 | 1.4 | 0.0 | 119 | 26 | 22.2 |
| **Total** | **86.0 ± 9.3** | **61.0 ± 13.3** | **11.1 ± 7.9** | **1.0 ± 1.9** | **2.9 ± 2.4** | **0.5 ± 1.0** | **131 ± 10** | **37 ± 10** | **28.2 ± 6.4** |

**Abbreviations:** **$TIR_{70-180}$**, Time In Range (70–180 mg/dL); **$TIR_{70-140}$**, Time In Range (70–140 mg/dL); **$TAR_{180}$**, Time Above Range (>180 mg/dL); **$TAR_{250}$**, Time Above Range (>250 mg/dL); **$TBR_{70}$**, Time Below Range (<70 mg/dL); **$TBR_{54}$**, Time Below Range (<54 mg/dL); **BG**, Blood Glucose; **SD**, Standard Deviation; **CV**, Coefficient of Variation

**Table S5. Individual swine closed-loop performance (mean ± SD) under AAPS control**. Performance metrics are reported for six sessions across two diabetic Yorkshire swine. All trials were conducted under an unannounced three-meal protocol with subject-specific tuning (basal rate, ISF, CR).

| | $TIR_{70\text{-}180}$ (%) | $TIR_{70\text{-}140}$ (%) | $TAR_{180}$ (%) | $TAR_{250}$ (%) | $TBR_{70}$ (%) | $TBR_{54}$ (%) | Mean BG (mg/dL) | SD (mg/dL) | CV (%) |
|---|---|---|---|---|---|---|---|---|---|
| **Subject 4 (Day 1)** | 69.8 | 50.5 | 29.5 | 7.7 | 0.7 | 0.0 | 156 | 49 | 31.6 |
| **Subject 4 (Day 2)** | 81.8 | 47.7 | 16.5 | 0.7 | 1.8 | 0.0 | 144 | 38 | 26.4 |
| **Subject 4 (Day 3)** | 42.6 | 6.2 | 57.4 | 8.8 | 0.0 | 0.0 | 192 | 39 | 20.2 |
| **Subject 5 (Day 1)** | 73.6 | 56.6 | 14.9 | 6.6 | 11.5 | 4.2 | 131 | 72 | 54.8 |
| **Subject 5 (Day 2)** | 66.0 | 60.4 | 11.1 | 4.5 | 22.9 | 9.7 | 112 | 56 | 49.5 |
| **Subject 5 (Day 3)** | 80.0 | 57.9 | 10.5 | 4.2 | 9.5 | 3.2 | 128 | 56 | 43.4 |
| **Total** | **69.0 ± 13.0** | **46.6 ± 18.5** | **23.3 ± 16.5** | **5.4 ± 2.7** | **7.7 ± 8.1** | **2.8 ± 3.5** | **144 ± 26** | **52 ± 12** | **37.6 ± 12.5** |

**Abbreviations:** $\mathbf{TIR_{70\text{-}180}}$, Time In Range (70–180 mg/dL); $\mathbf{TIR_{70\text{-}140}}$, Time In Range (70–140 mg/dL); $\mathbf{TAR_{180}}$, Time Above Range (>180 mg/dL); $\mathbf{TAR_{250}}$, Time Above Range (>250 mg/dL); $\mathbf{TBR_{70}}$, Time Below Range (<70 mg/dL); $\mathbf{TBR_{54}}$, Time Below Range (<54 mg/dL); **BG**, Blood Glucose; **SD**, Standard Deviation; **CV**, Coefficient of Variation

**Table S6. Individual swine AAPS control parameters.** Control profiles (basal rate, ISF, CR) for the two streptozotocin-induced diabetic Yorkshire swine (Subjects 4 and 5) engaged in the crossover benchmark study. These parameters were used exclusively for AAPS configuration.

| | Basal rate (U/h) | ISF (mg/dL/U) | CR (g/U) |
|---|---|---|---|
| **Subject 4** | 0.4 | 75 | 25 |
| **Subject 5** | 1.0 | 40 | 15 |

**Abbreviations: ISF,** Insulin Sensitivity Factor**; CR,** Carbohydrate Ratio
